# Tyrosine availability shapes *Staphylococcus aureus* nasal colonization and interactions with commensal communities

**DOI:** 10.1101/2025.05.06.651429

**Authors:** Laura Camus, Jessica Franz, David Gerlach, Anna Lange, Jeffrey John Power, Marcelo Navarro-Díaz, Johanna Rapp, Matthias Otto, Levin Joe Klages, Vladimir Starostin, Jens Mößner, Kevser Bilici, Soyoung Ham, Jörn Kalinowski, Largus T. Angenent, Hannes Link, Simon Heilbronner

**Author notes:** Lead contact. Simon Heilbronner. **Corresponding authors** Laura Camus, Simon Heilbronner.

## Abstract

Only certain individuals are nasally colonized by *Staphylococcus aureus*, making them more susceptible to developing endogenous infections. The role of the resident microbiota in modulating *S. aureus* nasal colonization remains largely unclear. Using a newly assembled nasal strain collection, we demonstrate that nasal commensals can either promote or inhibit *S. aureus* proliferation through strain- and community-specific effects *in vitro* and within gnotobiotic animals. We identified nutrient-dependent interactions as a key driver of these dynamics and found that *S. aureus* strains auxotrophic to tyrosine rely on amino acid-providing nasal communities to thrive in the nutrient-limited environment of the nose. Large-scale screenings revealed tyrosine dependency to affect 8% of nasal *S. aureus* strains and to impair *S. aureus* colonization. Our findings uncover a crucial factor in *S. aureus* physiology and its interactions as a commensal nasal species, paving the way for new decolonization strategies.

**Graphical abstract:** 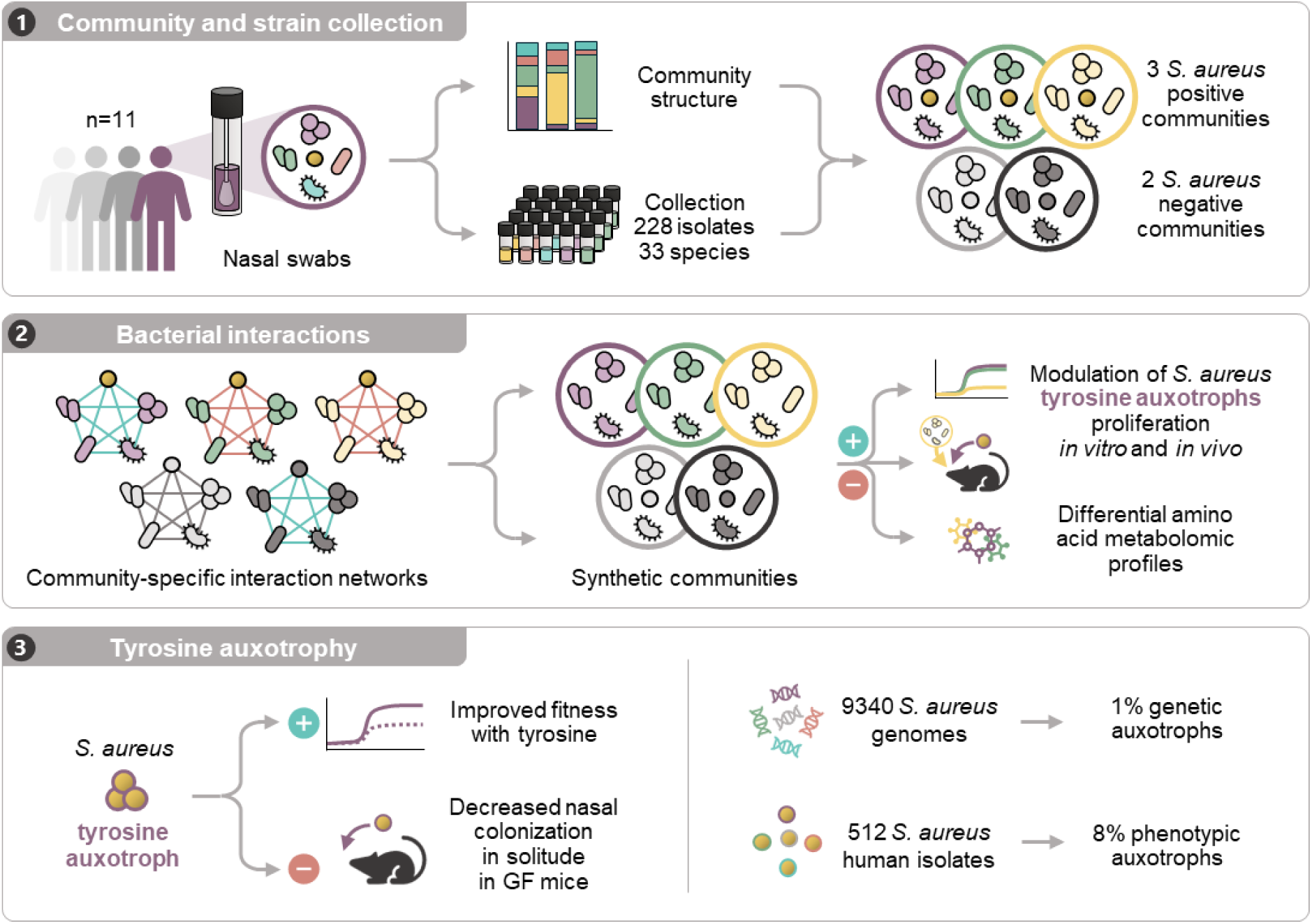

## Introduction

Host-associated microbial communities often act as reservoirs for opportunistic pathogens, thereby increasing the risk of infection. However, it has been recognized that certain commensal species can provide colonization resistance to invader species. Besides production of antibacterial and quorum-quenching molecules, nutritional competition has emerged as a crucial defense mechanism of the microbiota, particularly within the nutrient-rich surrounding of the human intestinal tract^1–3^. The nasal ecosystem can also harbor opportunistic pathogens, with *Staphylococcus aureus* constituting the greatest threat for human health due to its ability to cause a broad spectrum of infections^4^. Within the human population, individuals may be either permanently, transiently, or never colonized by *S. aureus,* and the underlying reasons are largely unclear. Only a few host, environmental, and bacterial factors have been shown to influence *S. aureus* nasal colonization and its behavior as a commensal species^4–6^. The role of resident microbial communities in shaping *S. aureus* proliferation within the nose is particularly complex, as both positive and negative interactions between *S. aureus* and individual nasal commensals have been reported^7, 8^.

In contrast to the gut ecosystem, the nose harbors a low density of both resources and bacteria, conditions that are expected to limit competitive behaviors^5, 9^. The nutrient scarcity within the nasal habitat is well documented, but has rarely been considered in previous interaction studies^7, 10–13^. Moreover, pairwise interactions alone do not necessarily predict the behavior of complex communities, as those can be influenced by high-order interactions^14^. In this context, the community composition, along with strain diversification and co-adaptation processes, were shown to shape the stability and colonization resistance of various microbiota^1, 2, 15–17^. Supporting the idea that similar dynamics may occur within nasal communities, recent studies highlighted that intra-species diversification influences the interaction of nasal commensals with *S. aureus*, notably through the production of antimicrobial or iron-scavenging molecules^13, 18^. However, the importance of strain-specific behaviors within complex nasal communities remains largely unexplored.

To address this gap, we established a nasal strain collection that captures intra-volunteer diversity at both the species- and strain-level. Using this collection, we show that interactions between *S. aureus* and nasal consortia are shaped by strain- and community-specific factors. Notably, we report that nasal *S. aureus* strains frequently exhibit tyrosine auxotrophy, making them reliant on amino acid-releasing communities to proliferate in the nutrient-poor environment of the nasal cavities.

## Results

### Establishment of the nasal strain collection LaCa

We established a novel nasal biobank to reconstruct nasal microbial communities at the strain level. We chose eleven healthy volunteers, determined the structure of their nasal microbiome using 16S rRNA gene amplicon sequencing, and isolated as many bacterial strains as possible from their anterior nares **(Fig. 1A)**. The analysis of the 16S rRNA gene amplicon sequencing showed distinct compositions, and ten out of eleven microbiomes could be assigned to a community state type (CST)^5^ number according to the dominance of *Corynebacterium* spp. (CST5, n=6), *Staphylococcus epidermidis* (CST3, n=2), *S. aureus* (CST1, n=1), and *Cutibacterium* spp. (CST4, n=1). One microbiome showed comparable abundances of *Corynebacterium* spp., *Cutibacterium* spp., and *Staphylococcus* spp., preventing a clear assignment to a CST **(Fig. 1B) (Fig. S1) (Tables S1 and S2)**. This analysis revealed a high abundance of *S. aureus* in communities B and C (27.1% and 75.8%, respectively), and a low abundance in A and D (0.2% and 1.6%, respectively). In each volunteer, 2 to 9 species (median: 4 species) accounted for the majority of the biomass (90% of sequenced 16S rRNA gene), while the remaining biomass was attributed to diverse species (18 to 51, median: 24 species) **(Table S2)**. These findings are in general agreement with previously published datasets^5, 19^.

**Fig. 1:**
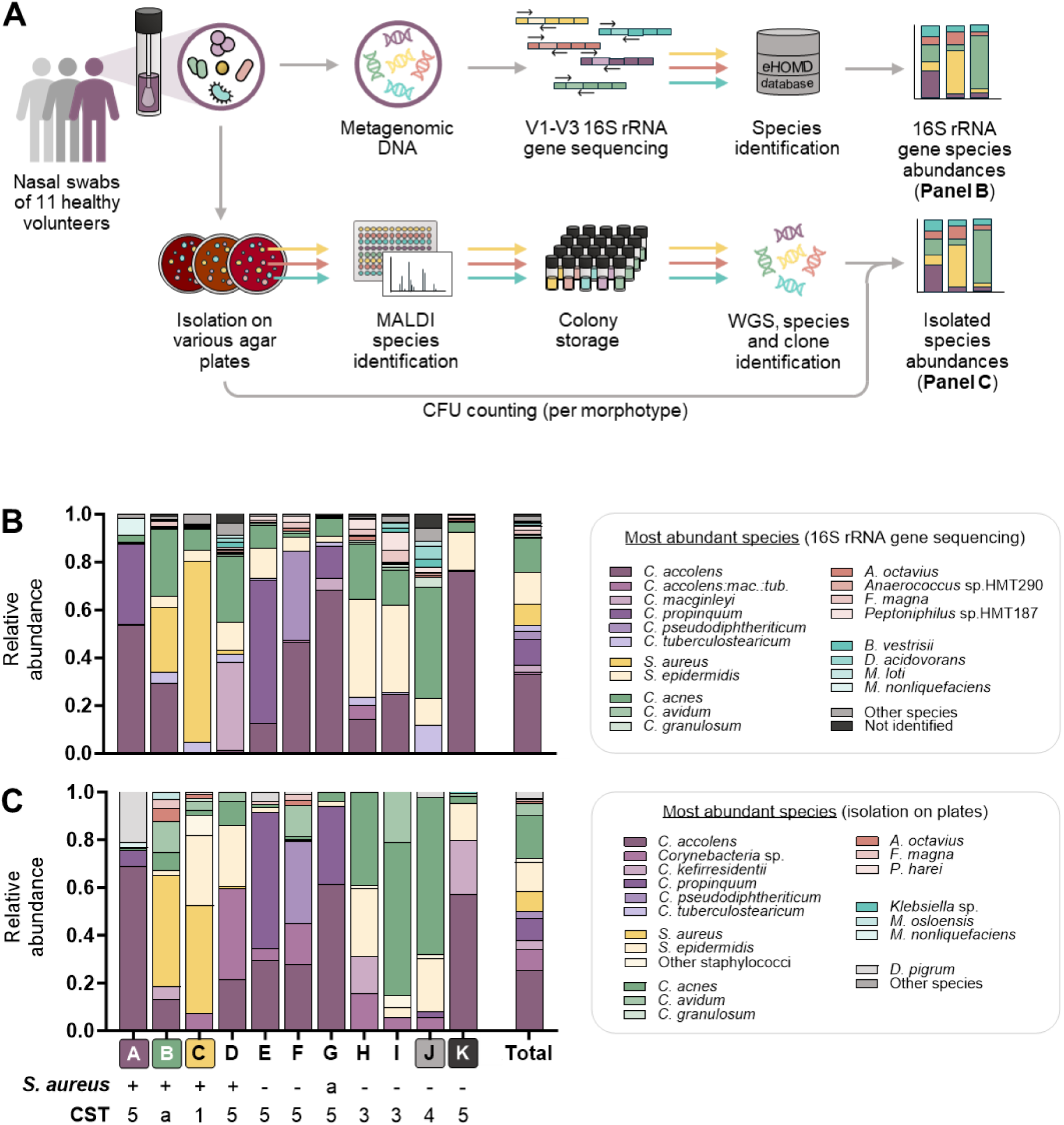
Species composition and structure of 11 nasal communities. **A.** Schematic methodology used to determine nasal community structures and establish the strain collection LaCa. **B, C.** Relative species abundance determined by 16S rRNA gene sequencing of nasal metagenomic DNA (**B**) or by isolation and enumeration of nasal commensals on plates (**C**), from the nasal microbiomes of eleven healthy volunteers (A-K). Each color designates one of the 20 most abundant species across all volunteers. Less abundant species were assigned to “Other species”. Species with a missing or unclear identification were assigned to “Not identified”. Below the bars, “+” indicates the volunteers in which *S. aureus* was detected by both techniques, and numbers correspond to the community state type (CST) assigned to the corresponding community. “a” refers to ambiguous *S. aureus* carriage (detected by only one method) or CST classification (balanced proportion of several species). Each bar displays the results of one replicate. See also Fig. S1, Tables S1, S2 and S3.

Using culturomics, 228 bacterial isolates (between 16 to 29 per volunteer) were selected based on colony morphology. The genomes of all strains were sequenced and analyzed **(Table S3).** After identification of clonal strains, the collection comprised 119 unique strains of 33 different species. Seven to 13 species were recovered per volunteer. Comparison to the 16S rRNA gene dataset showed that our approach successfully recovered the major species of the volunteers’ microbiomes **(Fig. 1C)**, with the exception of *Corynebacterium macginleyi* (abundant in Volunteer D) and strict anaerobes such as *Delftia acidovorans* or *Peptoniphilus* spp. (both abundant in Volunteers I and J). The intraspecies phylogeny of all isolates was assessed using the DSMZ Type Strain Genome Server^20^. In general, isolates of the same species were of clonal origin within an individual microbiome, but clonally unrelated across different microbiomes **(Table S3)**. However, we found a few exceptions. For instance, the volunteers B, E, F, J, and K were simultaneously colonized by two clonally distinct strains of *S. epidermidis.* Additionally, clonally identical strains of *Cutibacterium acnes* or *Corynebacterium tuberculostearicum/kefirresidentii* were found in 10 and 4 volunteers, respectively **(Table S3)**. These strains may have been transferred between the volunteers as they shared the same work environment.

### Nasal communities show different interaction networks

We speculated that strains colonizing the same human nose might show particular reciprocal support or inhibition. To investigate this, we selected volunteers A, B, C, J, and K showing different CST and carriage levels of *S. aureus* **(Fig. 1C)**, and quantified the binary interactions between co-isolated strains from each of these communities. We used an agar-based assay on synthetic nasal medium (SNM), which mimics the nutritional environment of the nasal cavity^10, 21^. Each strain was plated to form a lawn on SNM agar and incubated for six hours to foster growth and metabolic activity. All other strains of the community were then pinned on top of the lawn, or on SNM agar without a lawn as a control. After incubation, the area of arising colonies was assessed **(Fig. 2A)**.

**Fig. 2:**
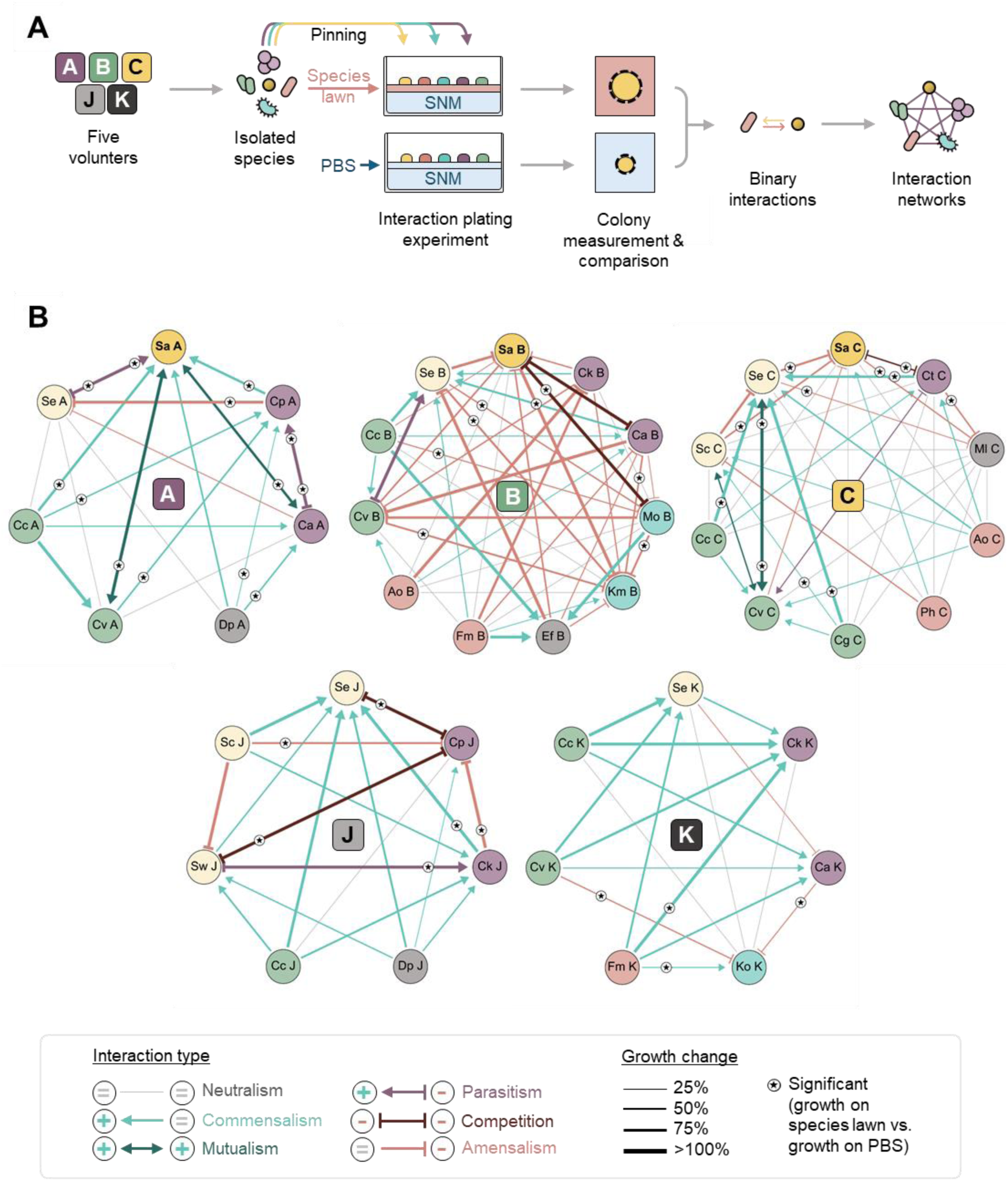
Binary interaction networks of the nasal communities A, B, C, J and K. **A.** Schematic methodology used to assess bacterial binary interactions within five communities. **B.** For each community, interactions were determined for every pair of nasal commensals on SNM agar and using colony area as a growth metric. Changes in the colony area were used to define the type and the strength of the interaction, as shown by the color, the shape and the thickness of the edges. Each edge is based on three biological replicates. The interactions inducing a significant change in the colony area of at least one of the two involved commensals are annotated with a star (*P*_adj_<0.05 two-way ANOVA with Dunnett’s correction). Nasal commensal species are indicated in the nodes. **Dark yellow:** *Staphylococcus aureus* (Sa). **Light yellow, other staphylococci:** *Staphylococcus epidermidis* (Se), *Staphylococcus capitis* (Sc), *Staphylococcus warneri* (Sw). **Green, *Cutibacteria*:** *Cutibacterium acnes* (Cc), *Cutibacterium avidum* (Cv), *Cutibacterium granulosum* (Cg). **Orange, anaerobic Gram-positives:** *Anaerococcus octavius* (Ao), *Finegoldia magna* (Fm), *Peptoniphilus harei* (Ph). **Blue, Gram-negatives:** *Klebsiella michiganensis* (Km), *Klebsiella oxytoca* (Ko), *Moraxella osloensis* (Mo). **Gray, other species:** *Dolosigranulum pigru*m (Dp), *Enterococcus faecalis* (Ef), *Micrococcus luteus* (Ml). **Purple, *Corynebacteria*:** *Corynebacterium accolens* (Ca), *Corynebacterium kefirresidentii* (Ck), *Corynebacterium propinquum* (Cp), *Corynebacterium tuberculostearicum* (Ct). See also Fig. S2.

Using this approach, we tested a total of 452 binary interactions, of which 266 interactions could not be evaluated as the involved strains failed to form detectable colonies on SNM agar or on the bacterial lawn. The remaining 186 interactions (28 to 55 per community) included both positive and negative interactions in every community, indicating that nasal commensals from the same microbiota can substantially impact each other’s proliferation **(Fig. 2B)**. The interaction network of each nasal community was found to be unique, and the proportion of positive and negative effects was variable across the different communities **(Fig. S2A)**. The communities A, J, and K were characterized by a high proportion of commensal interactions (47.4%, 61.1%, and 64.7%, respectively), while amensalism was less frequent (10.5%, 16.7%, and 17.7%, respectively). As reported previously, we found that *Corynebacterium propinquum* inhibited *S. epidermidis* proliferation^22^ in both communities A and J. In contrast, community B was dominated by amensal interactions (50%), while neutral interactions were most common in community C (36.8% of detected interactions) **(Fig. 2B) (Fig. S2A)**.

### Interactions of *S. aureus* with nasal commensals are strain- and community-specific

Based on the unique composition and interaction network of each nasal community, we hypothesized that distinct nasal communities may harbor a combination of traits that either promote or inhibit the proliferation of *S. aureus*. To assess this, we evaluated the binary interactions of *S. aureus* with the colonizing nasal commensals in more detail. The communities A, B, and C were *S. aureus*-positive based on 16S rRNA gene analysis and culturomics **(Fig. 1B** and **C)**. The corresponding *S. aureus* strains were designated Sa A, Sa B, and Sa C, respectively. The only additional trait shared by communities A, B, and C was the presence of distinct strains of *S. epidermidis* and *C. avidum* **(Table S3)**. Interestingly, the interactions of *S. aureus* with other species differed substantially between the three communities. The proliferation of Sa A was enhanced by most members of the community A, including *S. epidermidis* and *C. avidum*. In contrast, we found Sa B and Sa C to be either inhibited or not affected by strains of community B and C, respectively **(Fig. 2B) (Fig. S2B and C)**. Similarly, the *S. epidermidis* and *C. avidum* isolates showed different interaction patterns with each of the three *S. aureus* strains **(Fig. S2C)**.

In complex communities, the outcome of pairwise relationships can be influenced by the presence of other species, which is a phenomenon known as high-order interactions^14^. To assess this, the *S. aureus* strains Sa A, Sa B, Sa C, and the reference strain Sa Newman were grown on SNM agar in solitude or within synthetic communities **(Fig. 3A** and **B)**. The SynComs were assembled to represent the species and strain composition of the microbiomes from volunteers carrying (A, B, and C) or not carrying *S. aureus* (J and K). When we assessed the effects of the SynComs A, B, and C, we found the reference strain Sa Newman to be significantly supported by SynCom B. The two other SynComs had a positive but non-significant impact on Sa Newman’s growth in synthetic culture **(Fig. 3B) (Fig. S3)**. Sa B and Sa C profited from the SynComs B and C, respectively, but were hardly affected by the other ones. In contrast, Sa A proliferation was increased in the presence of communities A and B, but limited by community C **(Fig. 3B) (Fig. S3)**. The SynComs J and K showed a general ability to improve *S. aureus* growth, challenging the idea that communities derived from *S. aureus*-negative volunteers might exert general inhibitory effects on the pathogen **(Fig. 3B) (Fig. S3)**. To assess whether contact-dependent mechanisms or secreted products mediated the effects observed in synthetic culture, we cultivated *S. aureus* Newman, Sa A, Sa B, and Sa C in spent culture supernatants of the SynComs. The supernatant of the SynComs J and K significantly enhanced the growth of all *S. aureus* strains in a similar pattern to the one observed in the synthetic culture experiments **(Fig. S4A and B)**. Importantly, Sa A was also strongly supported by the supernatants of the SynComs A and B, whilst not growing in the spent medium of the SynCom C **(Fig. S4C)**, suggesting that this strain is influenced by community-secreted products.

**Fig. 3:**
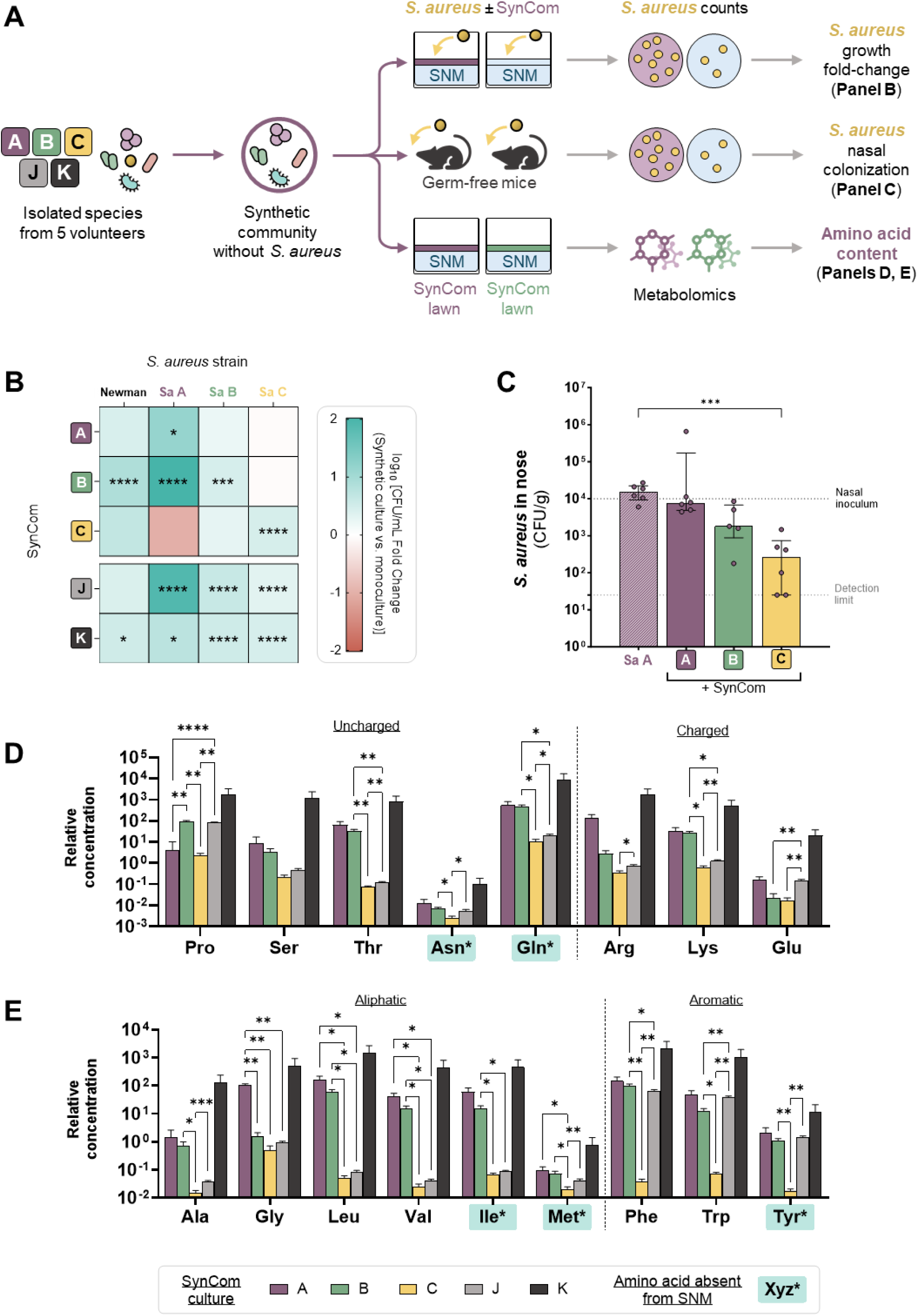
Impact of SynComs on S. aureus proliferation and amino acid composition of SNM. **A.** Schematic methodology used to study synthetic communities *in vitro* and *in vivo*. **B.** Heatmap depicting the growth fold-change induced by synthetic culture with five different SynComs (A, B, C, J, K) on SNM agar, for four *S. aureus* strains. Sa A, Sa B and Sa C originated from volunteers A, B, and C, respectively. Positive fold-changes (blue) and negative fold-changes (red) indicate increases or decreases in *S. aureus* cell numbers in synthetic culture relative to the monoculture, respectively. Each fold-change is based on four biological replicates. Significant changes are marked with stars (\**P*<0.05, \*\**P*<0.01, \*\*\**P*<0.001, \*\*\*\**P*<0.0001, 2-way ANOVA). See also Fig. S3. **C.** Barplots depicting the bacterial counts of *S. aureus* Sa A recovered from the noses of gnotobiotic mice, seven days after nasal inoculation in isolation or in the presence of three different SynComs. Bars represent the median ± interquartile range from 5 to 6 animals per group. Dashed lines represent the nasal inoculum and the detection limit. Statistical significance is indicated by \*\**P*_adj_<0.01 (Kruskal-Wallis with Dunn’s correction). **D, E.** Bar plots showing the concentration of polar (**D**) and nonpolar (**E**) amino acids detected in the cell extract of five different SynComs (A, B, C, J, K) cultivated on SNM agar. Cysteine and histidine were not assessed due to analytical limitations, and aspartic acid was not detected in the samples. Amino acids marked with a star and highlighted in blue are not present in the original composition of SNM. LC-MS data of SynCom samples were normalized to the growth of the respective community, as measured by optical density of the cell extract. Bars represent the mean relative concentration + SD from four biological replicates, represented as log_10_-transformed values. Statistical significance is indicated by \**P*_adj_<0.05, \*\**P*_adj_<0.01, \*\*\**P*_adj_<0.001, \*\*\*\**P*_adj_<0.0001 (Two-way ANOVA with Tukey’s correction). See also Fig. S5 and Table S4.

To determine whether these effects are relevant *in vivo*, we developed a nasal colonization model using germ-free mice and tested the impact of SynComs A, B, and C on the proliferation of Sa A **(Fig. 3A** and **C)**. In the absence of nasal commensals, 1,5×10^4^ CFU/g of Sa A could be recovered from the mice’s noses after seven days of colonization. Importantly, the nasal bacterial loads of Sa A were maintained in the presence of SynCom A and B but were reduced by a 2-log factor in the presence of SynCom C, largely reflecting the pattern observed during *in vitro* synthetic cultures **(Fig. 3B** and **C)**. Taken together, these results show that the impact of nasal commensals and complex consortia on *S. aureus* proliferation is highly strain-specific, preventing the identification of a general pattern. Nasal communities could either support or inhibit *S. aureus,* depending on the strain, with Sa A being the most affected isolate in our collection.

### Communities affect the amino acid composition of the nasal environment differently

Amino acids are essential for bacterial growth. Nasal secretions are devoid of asparagine, aspartate, glutamine, isoleucine, methionine, and tyrosine, and these are, therefore, not present in SNM^10^. We speculated that changes in amino acid availability might explain the effects of nasal communities on *S. aureus*. To investigate this, we compared the amino acid profiles of SynComs culture extracts **(Fig. 3A, D**, and **E) (Table S4)**. Principal component analyses showed each community to have a distinct impact on amino acid availability **(Fig. S5)**. Five of the six amino acids that are missing from SNM were detected in all SynCom cultures, suggesting that nasal communities can enhance the nutrient diversity in the medium. SynCom K presented the highest amino acid concentrations, whereas SynCom C and J showed generally lower levels **(Fig. 3D** and **E)**. Additionally, SynCom C was characterized by reduced concentrations of the aromatic amino acids tyrosine, tryptophan, and phenylalanine compared to all other communities, suggesting that it may impose a particularly strong nutritional stress on *S. aureus* **(Fig. 3E)**.

### Tyrosine auxotrophy drives *S. aureus* interaction with nasal communities

We hypothesized that the nutritional stress caused by limited amino acid availability in SynCom C underlies its growth-restricting effect on Sa A. Only Sa A showed a growth defect in SNM but proliferated in rich medium **(Fig. S6)**. Supplementing SNM with the six lacking amino acids in a mixture or individually showed tyrosine to be necessary and sufficient to restore the growth of Sa A in a dose-dependent manner **(Fig. S7)**. This suggested Sa A to be auxotrophic for tyrosine.

We sequenced the genome of Sa A and identified the coding sequence of *tyrA* to be disrupted by a frameshift mutation (T insertion in position 817/1093), resulting in a premature stop codon **(Fig. S8)**. The tyrosine biosynthesis pathway of Staphylococci has not yet been characterized in detail. However, the prephenate dehydrogenase TyrA is known to be essential for tyrosine biosynthesis in *Bacillus* spp., *Escherichia coli*, and other species^23, 24^. Genomic reversion of the frameshift mutation in Sa A enabled growth in SNM **(Fig. 4A** and **B)**. The growth-enhancing effect of the SynCom A on Sa A was abrogated when the frameshift in *tyrA* was removed, indicating that this mutation caused dependency of the strain on tyrosine released by the community **(Fig. 4C)**. Introduction of Δ*tyrA* mutations in *S. aureus* Newman and USA300 JE2 induced tyrosine auxotrophy and community-dependency, as observed for Sa A **(Fig. S9)**. Altogether, these findings show *tyrA* deficiency to cause tyrosine auxotrophy in *S. aureus,* making it reliant on communities that provide this amino acid in nutritionally poor environments.

**Fig. 4:**
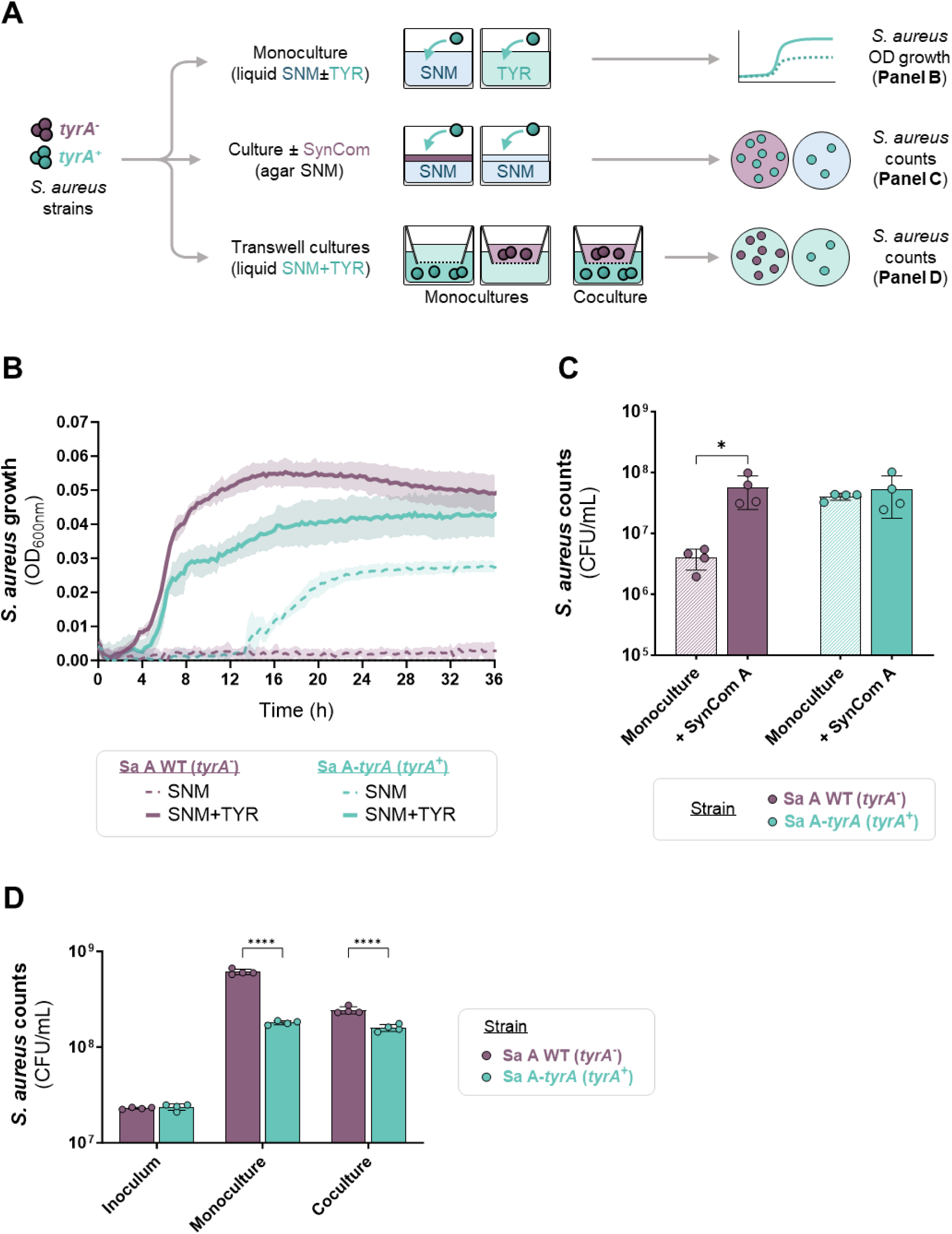
Effect of tyrA impairment on the proliferation of *S. aureus* Sa A in SNM. **For all panels:** the nasal *S. aureus* strain with an impaired *tyrA* gene (Sa A WT) is referred as *tyrA*^-^ and shown in purple, while the strain with the restored *tyrA* gene (Sa A-*tyrA*) is named *tyrA*^+^ and shown in blue. Lines or bars represent mean ± SD of four biological replicates. Statistical significance is indicated by \**P*_adj_<0.05, \*\*\*\**P*_adj_<0.0001 (Two-way ANOVA with Šídák’s correction). A. Schematic methodology used to study the behavior of *S. aureus tyrA*^-^ and *tyrA*^+^ strains. B. OD_600nm_ growth kinetics in SNM and SNM + tyrosine (TYR). See also Fig. S9A and C. C. Barplots showing the bacterial counts of *S. aureus* cultivated on SNM agar in isolation or in the presence of the SynCom A over 3 days. See also Fig. S9B and D. D. Barplots showing the bacterial counts of *S. aureus tyrA*^+^ and *tyrA*^-^ in the inoculum, cultivated in SNM + tyrosine either in isolation or cocultured with each other over 24 hours using the transwell system. See also Fig. S10.

### Tyrosine auxotrophic strains exhibit a fitness advantage in the presence of tyrosine

Auxotrophs are common in bacterial communities and impact their diversity and stability, often through cross-feeding between individuals with different metabolic requirements^25, 26^. Reduction of metabolic labour offers auxotrophs a fitness advantage when they acquire the missing amino acid from an external source^27, 28^. To assess this in *S. aureus*, we conducted mono- and cocultures of Sa A WT strain (*tyrA*^-^) and *tyrA-*restored mutant (*tyrA*^+^) in SNM + tyrosine using a transwell system that allows nutrient exchange between separately growing strains **(Fig. 4A** and **D)**. In the presence of tyrosine, Sa A WT reached higher cell densities than the *tyrA*-restored mutant, both in monoculture and when strains competed for the shared nutrients **(Fig. 4D) (Fig. S10A)**. Similar effects were observed in the Sa Newman and Sa JE2 background **(Fig. S10)**, suggesting that tyrosine auxotrophy provides a general selective advantage when the amino acid is supplied by external sources.

### Nasal *S. aureus* strains exhibit various amino acid requirements

We examined the frequency of frame-shift mutations in the *tyrA* gene across 9340 *S. aureus* genomes and found the gene to be potentially disrupted in ≈1% of cases (n= 98 genomes). However, phenotypic auxotrophy seemed more informative because the altered primary structure of TyrA, along with mutations in other genes of the pathway or in its regulatory network, might also cause auxotrophy. We gathered a biobank of 512 *S. aureus* human isolates and assessed their growth on standard SNM and SNM supplemented with tyrosine **(Fig. 5A) (Fig. S11A, B and C)**. Forty-two isolates (8.2%) failed to proliferate on SNM. Supplementation of the medium with tyrosine rescued the growth of five of them, and three additional isolates formed colonies only in the presence of the six amino acids missing from SNM **(Fig. 5B** and **C)**. However, among strains that were able to grow in plain SNM, we found 47 (9.2% of total strains) whose proliferation was significantly stimulated by supplementation with tyrosine or the six amino acid mixture **(Fig. 5B** and **C) (Fig. S11D and E)**. When focusing on strains with well-defined isolation sites, we found a higher frequency of amino acid dependency and growth defects in SNM in nasal isolates compared to those from infection sites **(Fig. 5D)**. These findings suggest that nasal *S. aureus* strains do frequently exhibit auxotrophies to various amino acids, which can partially or completely hinder their proliferation in SNM.

**Fig. 5:**
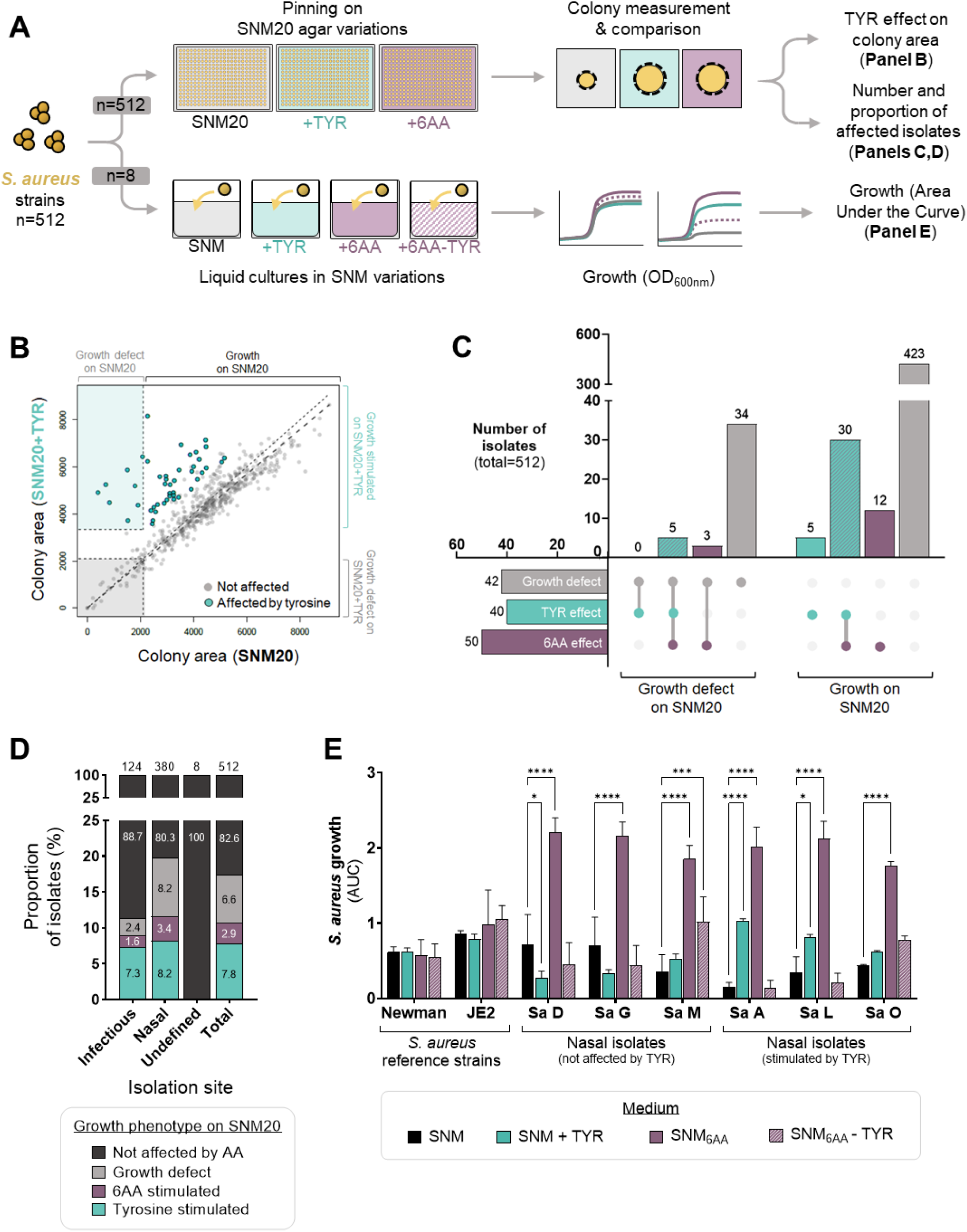
Phenotypic tyrosine dependency of *S. aureus* isolates. **A.** Schematic methodology used to identify phenotypic auxotrophies in *S. aureus* strains. See also Fig. S11A, B and C. **B.** Dotplot depicting the effect of tyrosine (TYR) supplementation on the growth on SNM20 of a collection of *S. aureus* isolates (n= 580 depicted colonies; n=525 unique isolates). Colored zones denote thresholds that categorize isolates with growth defects on SNM20 that can either be restored (blue zone) or not (grey zone) by tyrosine supplementation. The diagonal dashed lines represent the linear regression corresponding to a ratio of one (thin line) or the actual linear regression (thick line), showing that most *S. aureus* were not affected by tyrosine supplementation. Isolates showing a significant growth improvement in the presence of tyrosine are highlighted in blue (*P*_adj_<0.05, t-test with Bonferroni correction). Each dot represents the colony area from three biological replicates. See also Fig. S11. **C.** Upset diagram showing responses to amino acid supplementation across 512 *S. aureus* isolates. Isolates from human origin and unique clonality were selected from the screening shown in Panel A. Horizontal bars represent the number of isolates exhibiting a growth defect on SNM20 agar, or a growth stimulation by supplementation with either the six amino acid mix (6AA) or tyrosine (TYR) alone. Vertical bars indicate the number of isolates showing each growth phenotype, sorted by their basal ability to form colonies on SNM20 without amino acids. **D.** Bar plots showing the proportion of *S. aureus* human isolates not affected, showing a growth defect on SNM20 or being stimulated by tyrosine or the six amino acid mix, according to their isolation site. Numbers within bars indicate the percentages of strains with the observed phenotype. Numbers above bars indicate the number of tested strains. **E.** Bar plots showing the proliferation of 8 *S. aureus* strains in SNM or in SNM supplemented with tyrosine, with a mixture of the six amino acids missing from SNM, or with the same mixture from which tyrosine was removed (6AA-TYR). 100µM of each amino acid was used. Each bar displays the mean AUC + SD from three biological replicates of growth kinetics. Statistical significance is indicated by \**P*_adj_<0.05, \*\*\**P*_adj_<0.001, \*\*\*\**P*_adj_<0.0001 (Two-way ANOVA with Dunnett’s correction).

To further validate this, we selected five nasal isolates that responded differently to tyrosine supplementation in our initial screen and cultured them in liquid SNM with various amino acid combinations **(Fig. 5A)**. Tyrosine supplementation enhanced the proliferation of Sa O and Sa L, but not that of Sa D, Sa G, and Sa M. The growth of these isolates was rescued in the presence of the six amino mix, but only when tyrosine was included **(Fig. 5E) (Fig. 6A, B and C)**. This suggests that tyrosine is necessary but not sufficient for these isolates to grow in SNM, likely due to multiple amino acid auxotrophies. Proliferation of Sa G, Sa L, and Sa O, in SNM was enhanced by SynCom supernatants or living cells, largely reflecting the behavior of Sa A **(Fig. 6D** and **E)**. Genome analysis showed the *tyrA* gene of Sa G to be disrupted by a frame-shift mutation (T deletion in position 325/1091) **(Fig. S8)**. In contrast, no evident detrimental *tyrA* mutations were identified in the remaining isolates, suggesting that other mutations or regulatory mechanisms may underlie their dependency on amino acid supplementation. Collectively, these results highlight that nasal *S. aureus* strains have diverse amino acid requirements, with tyrosine metabolism and sharing being crucial for *S. aureus* proliferation within nasal communities.

**Fig. 6:**
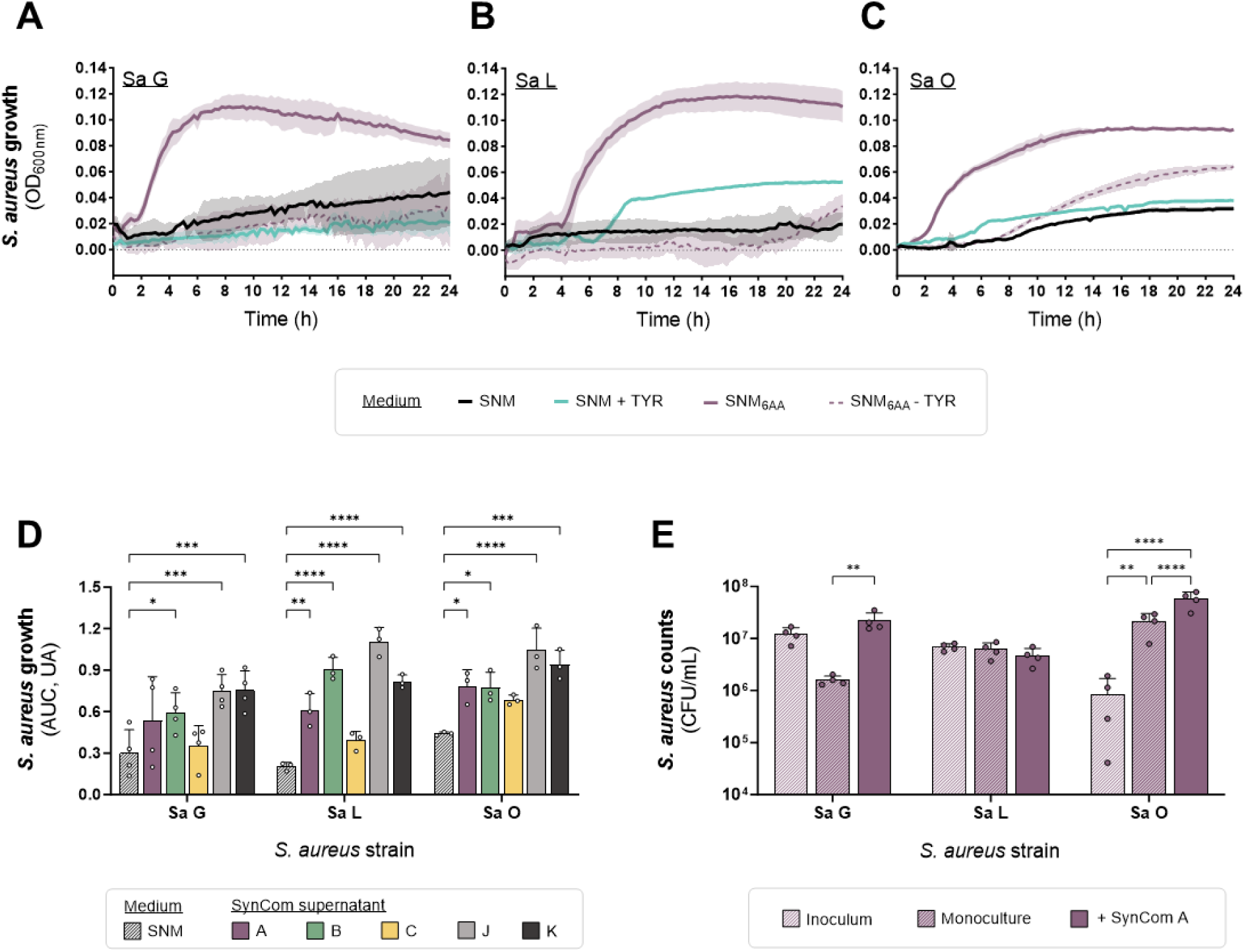
Growth behavior of nasal S. aureus strains with amino acid dependencies. A, B, **C.** Proliferation of Sa G (**A**), Sa L (**B**) or Sa O (**C**) in SNM or in SNM supplemented with tyrosine (TYR), with a mixture of the six amino acids missing from SNM (6AA), or with the same mixture from which tyrosine was removed (6AA-TYR). 100µM of each amino acid was used. Lines represent the mean ± SD of three biological replicates of OD_600nm_ measurements, taken every 15 minutes over 24 hours. **D.** Proliferation of Sa G, Sa L or Sa O in SNM or in the presence of 50% concentrated spent culture supernatant of the SynComs A, B, C, J and K. Each bar displays the mean AUC + SD from three to four biological replicates of growth kinetics. Statistical significance is indicated by \**P*_adj_<0.05, \*\**P*_adj_<0.01, \*\*\**P*_adj_<0.001, \*\*\*\**P*_adj_<0.0001 (Two-way ANOVA with Dunnett’s correction) **E.** Barplots depicting the bacterial counts of Sa G, Sa L or Sa O in the inoculum, and after incubation for 3 days in monoculture and in presence of the SynCom A on SNM agar. Bars represent the mean + SD of four biological replicates. Statistical significance is indicated by \*\**P*_adj_<0.01, \*\*\*\**P*_adj_<0.0001 (Two-way ANOVA with Šídák’s correction).

### Tyrosine auxotrophy impairs the ability of *S. aureus* to colonize the nasal cavity *in vivo*

To further validate the importance of amino acid biosynthesis for proliferation *in vivo*, we compared the ability of *S. aureus* Newman WT and Δ*tyrA* to colonize the nostrils of gnotobiotic animals. Tyrosine auxotrophy reduced *S. aureus* nasal cell counts by two orders of magnitude **(Fig. 7A)**. Interestingly, these differences were mitigated when *S. aureus* was enumerated in murine feces **(Fig. 7B)**, suggesting that tyrosine auxotrophy specifically impairs *S. aureus* colonization of the nose in the absence of other commensals.

**Fig. 7:**
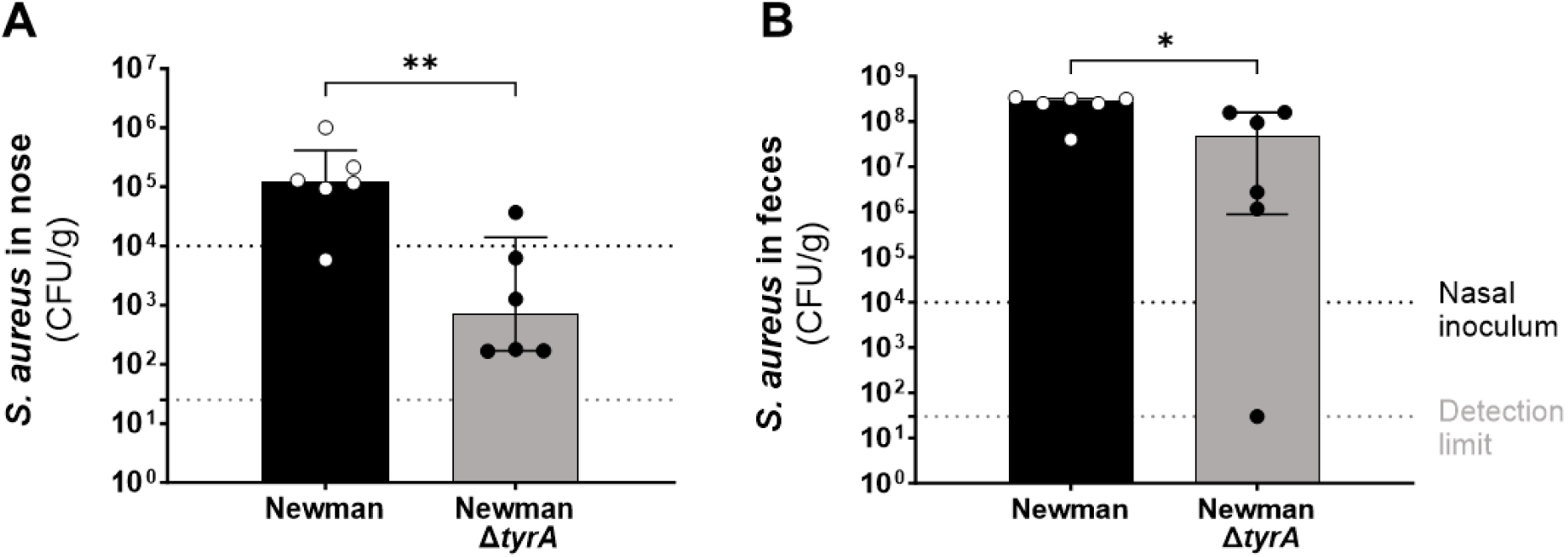
Effect of *tyrA* deficiency on the ability of *S. aureus* to colonize the nasal cavities and the intestine of gnotobiotic mice. Barplots depicting the bacterial counts of *S. aureus* Newman WT and Δ*tyrA*, recovered from the noses (**A**) or feces (**B**) of gnotobiotic mice seven days after nasal inoculation. Bars represent the median ± interquartile range from 5 to 6 animals per group. Dashed lines represent the nasal inoculum and the detection limit. Statistical significance is indicated by \**P*_adj_<0.05, \*\**P*_adj_<0.01 (Mann-Whitney test).

## Discussion

We identify commensalism as the most common type of interaction among nasal bacteria, dominating four out of five interaction networks in our study. In contrast, competition is described as the dominant type of interaction in gut and soil microbiota, where they are thought to play a greater role in stabilizing community structures than positive interactions^9, 29^. This difference may stem from the distinct nutritional conditions of the nasal cavities, which are characterized by a lower resource diversity and density^9, 10^. This finding suggests that bacterial interactions – and, by consequence, colonization resistance – are driven by different processes in the nose compared to other ecosystems. Several studies have evidenced the existence of amensalistic interactions, notably through the direct killing of *S. aureus* by nasal commensals producing antimicrobial molecules^7, 12^. In contrast, competitive exclusion within the nasal microbiome has so far been described only in the context of siderophore-mediated iron acquisition^13, 22, 30^. By employing an unbiased and high-throughput approach to analyze interactions in a medium mimicking the nutritional environment of the nose^10^, we discovered that nasal communities could also support *S. aureus* proliferation by providing essential nutrients such as amino acids.

While less frequent than competitive interactions, cross-feeding between auxotrophic strains is commonly observed within microbial communities, contributing to their diversity and stability^25, 31, 32^. We found that 8.2% of nasal *S. aureus* strains were stimulated by tyrosine supplementation and, therefore, likely to be supported by amino acid-releasing nasal communities. However, the degree of tyrosine dependency varied across strains, suggesting that this phenotype may be selected when tyrosine is supplied by external sources^27, 28^. The enhanced fitness of Sa A in the presence of tyrosine, along with its isolation from a community that releases high levels of this amino acid, further supports this hypothesis. Interestingly, metagenomic analyses revealed that Sa A and Sa G were present at low relative abundance within their respective communities. Whether the tyrosine auxotrophy of these isolates limits their competitive behavior, and if this might contribute to a long-term stability of the consortium^25^, remains to be explored. It is also possible that auxotrophic *S. aureus* strains preferentially colonize, or might even be selected by nasal communities that provide the missing nutrient. This could be investigated through longitudinal screenings for auxotrophic *S. aureus* strains, combined with advanced metabolomics techniques to detect tyrosine and/or other relevant metabolites in the nose. Indeed, in light of our results and of those from other studies^31^, it seems likely that *S. aureus* can benefit from additional public goods provided by resident commensals, including other amino acids or vitamins. In turn, *S. aureus* can also provide essential resources, such as siderophores, and support other nasal species and communities^13^.

The impact of public goods and related auxotrophies on the ability of *S. aureus* to colonize ecosystems beyond the nose remains an open research question. Our *in vivo* experiments demonstrated that tyrosine auxotrophy affects nasal, but not intestinal colonization, possibly due to the greater resource availability in the gut^33^. Additionally, *tyrA* transposon mutants did not show reduced potential for causing infections^34–37^ or proliferating in the blood or cystic-fibrosis lung sputum^38, 39^. This suggests that tyrosine is sufficiently available in many biological habitats of *S. aureus,* and that auxotrophy may actually be selected during nasal colonization in the context of certain bacterial communities. Consistent with this, a recent genomic study found the metabolic pathways of tryptophan, phenylalanine, and tyrosine to be more frequently affected by non-synonymous mutations in *S. aureus* isolates from colonization compared to those from infections^6^. If these mutations have functional consequences remains elusive. Dissecting the tyrosine biosynthetic pathway and its regulation in *S. aureus* would provide deeper insights into its role in shaping the bacterium’s physiology and microbial interactions.

In summary, our study shows that the ability of *S. aureus* to colonize the nasal ecosystem and to interact with its bacterial members is highly dependent on community- and strain-specific features. Among these, we identified amino acid metabolism as a key component of the colonization mechanisms of *S. aureus*, paving the way for finding new strategies to understand and control its persistence in the nose.

## Methods

### Bacterial strains

The bacterial strains used in this study are listed in **Table S3** and **Table S5**. The nasal strains from the LaCa collection were isolated from the nose of eleven healthy adults. The sample collection procedures were approved by the clinical ethics committee of the University of Tübingen (No. 109/2009 BO2) and informed written consent was obtained from the volunteers. The complete protocols for strain isolation and identification are provided in the **Supplementary Methods**. Briefly, one nasal swab per volunteer was used for strain isolation, as previously described^19^. Species were identified using matrix-assisted laser desorption/ionization-time-of-flight mass spectrometry as described elsewhere^40^. Two to four isolates per species and per volunteer were frozen to constitute the collection LaCa **(Table S3)**. Species identification was confirmed by full genome sequencing **(Supplementary Methods)**.

The clean deletion of *tyrA* in *S. aureus* Newman and USA300 JE2, as well as the restoration of this same gene in Sa A were generated via allelic exchange with the thermosensitive plasmids pIMAY:Δ*tyrA* and pIMAY:*tyrA*(T)^41, 42^ **(Table S5 and S6)**. Briefly, 500 bp flanking regions upstream or downstream of the *tyrA* gene were amplified from chromosomal DNA using A/B and C/D primers with overlapping sequences. A 1 kb deletion fragment with an ATG-TAA scar was created using spliced overlap extension PCR (SOE-PCR). Similarly, the reversion cassette to remove the frameshift in Sa A was created by removing the inserted nucleotide in primers B and C. Constructs were then cloned into pIMAY by restriction digestion and used to transform *S. aureus* target strains. Mutants were created using allelic exchange as previously described^41, 42^.

### Culture conditions

For all experiments, bacterial precultures were prepared in BHI (Roth), BHI + 0.2% Tween 80 (Sigma) or TSB (Oxoid) and incubated under conditions specific to each species, as detailed in **Table S7**. For strict anaerobes with limited growth in liquid broth, bacterial suspensions were prepared by harvesting colonies from fresh agar plates. Bacterial cells from the preculture or from the colonies were then washed twice in sterile PBS prior to concentration adjustment. Subsequent experiments were conducted in variations of Synthetic Nasal Medium (SNM, SNM3, SNM20), prepared as described previously^10^. Depending on the assay, the medium was supplemented with the six amino acids missing from SNM either individually or as a mixture (100 µM each). These amino acids included: tyrosine, methionine, glutamine, asparagine, aspartate (AppliChem) and isoleucine (Roth). In general, growth kinetics over 24 h or 36 h were conducted in 48-well plates at 37°C with continuous orbital shaking. OD_600nm_ was measured every 15 min using an Epoch plate reader (BioTek).

### Binary interactions between nasal commensals

Pairwise interactions between nasals commensals were assessed using the most abundant isolate of each species derived from the volunteer of interest, based on bacterial counts obtained during the strain isolation process **(Supplementary Methods) (Table S3)**. Interaction assays were conducted as described previously^21^. Briefly, the suspension of a background species, or PBS as control, was spread as a lawn on SNM agar containing bromocresol blue as a stain in rectangle plates (Singer). The plate was incubated at 37°C for six hours under appropriate oxygenation conditions to foster growth and metabolism of the background species **(Table S7)**. All strains to be tested were then pinned manually on top of the lawn using 96-pins pads (Singer). Plates were incubated again for seven days at 37°C in aerobiosis and imaged under standardized conditions, using a light pad (Gaomon) and a 14.1 Mpx camera (Panasonic DMC-FZ100). We developed and used the software “BactoVision” to measure the area of arising colonies. This tool ensures standardized measurement of colonies across plates, incorporating semi-automated image enhancement features and colony detection (https://github.com/mlcolab/bactovision). The impact of the background species on each pinned species was quantified by a growth fold change, calculated by dividing the colony area measured on the background species lawn by the one obtained on the PBS lawn. Interactions were categorized as detailed in **Fig. 2B**, then plotted as network diagrams.

### Effect of SynComs on *S. aureus* proliferation

Synthetic communities (SynComs) were assembled using the same isolates used in the binary interaction experiments. For the SynComs B and K, the species *K. oxytoca* and *K. michiganensis* were excluded due to their overgrowth of other species and their absence in the results of the 16S rRNA gene amplicon sequencing **(Table S1)**. Following preculture in rich medium, each isolate was resuspended in PBS-glycerol to OD_600nm_=5, after which the suspensions were combined in an equal ratio. Aliquots of the resulting SynCom were stored at -80°C until further use. The absence of contaminants and the presence of all expected species were verified by plating and 16S rRNA gene amplicon sequencing **(Fig. S12)**. For synthetic culture assays, OD_600nm_=1 suspensions were prepared from the frozen SynCom and a fresh preculture of *S. aureus*. Twenty microliters of *S. aureus*, the SynCom or both (1:1 ratio) were evenly distributed onto SNM agar in 12-well plates and incubated at 37°C for three days. To determine bacterial concentrations, the agar was flooded with 1 mL PBS to harvest cells and enumerate them on TSA+Blood or Chromagar *S. aureus*^43^. For *S. aureus* exposure to secreted metabolites, each SynCom was cultivated in liquid SNM at 37°C with agitation for 48 h, after which cultures were filtered. Growth kinetics of *S. aureus* in liquid SNM in presence or in absence of 50% spent supernatant of the SynCom were then conducted as described in “Culture conditions”.

### Metabolome of SynComs

Amino acid quantification in cell extracts was conducted by cultivating SynComs in monoculture for 3 days on SNM agar at 37°C, using the 12-well plate format as described above. After incubation, 1 mL of 80% methanol was added to each well, incubated for 20 min and recovered for OD_600 nm_ measurement. The methanol extract was centrifuged at 12,000 rpm for 10 min and the supernatant was stored at -80°C until analysis. Targeted metabolomics was performed via isotope ratio LC-MS/MS on an Agilent TQ 6495. For chromatographic separation, an InfinityLab Poroshell 120 HILIC-Z 2.7 µm, 2.1x150mm (Agilent Technologies) was used. Chromatographic parameters were set as described in Agilent application note 5994-5949. TQ settings and multiple reaction monitoring were obtained from Guder and colleagues^44^. For analysis, the LC-MS metabolite ratios of the SynComs were normalized to their respective OD600nm **(Table S4)**.

### Cocultivation of prototrophic and auxotrophic *S. aureus* strains

The transwell system (Corning) was used to perform mono- and cocultures of *S. aureus* strains with a functional (Newman WT, USA300 JE2 WT, Sa A-*tyrA*) or an impaired (Newman Δ*tyrA,* USA300 JE2 Δ*tyrA*, Sa A WT) *tyrA* gene **(Table S5)**. Bacterial suspensions with an OD_600 nm_ 0.015 were prepared in SNM + 100 µM tyrosine and inoculated either into the insert (500 µL, *tyrA*^-^ strains) or in the bottom part (1,5 mL, *tyrA*^+^ strains) of a 12-well transwell plate. For monocultures, the bacterial suspension of one partner was replaced by sterile SNM + tyrosine. The plate was incubated for 24 h at 37°C with agitation. Bacterial cell counts were determined for the insert and bottom parts of each well by plating on TSA.

### *In vitro* identification of tyrosine auxotrophs

A biobank was constituted with 512 *S. aureus* human isolates from the collections of J. Larsen^45^ (JeLa, n=8), J. Rudkin (JuRu, n=277), students from Munich (MuSt, n=202), Universitätsklinikum Tübingen (UKT, n=19), and this study (LaCa, n=6). This biobank included isolates of nasal (n=380), infectious (n=124) or undefined (n=8) origins. To validate the method, a set of control and reference strains were also included in the pinning assay **(Fig. S11A and B) (Table S5)**.

A visualization of the screening procedure is depicted in **Fig. 5A**. Bacterial suspensions of all strains were frozen in cryoprotective medium at -80°C in microtiter plates until use. Using the PM1 robotic colony replicator (S&P Robotics)^46^, the biobank was arranged into the 384 well microtiter format on rectangle plates (VWR) containing TSA and cultivated overnight at 37°C for recovery. For the high-throughput detection of tyrosine dependencies, a 20x-concentrated version of the SNM (SNM20)^10^ was used to ensure robust growth and subsequent colony detection. The biobank was transferred on SNM20 agar either unmodified or supplemented with tyrosine alone or a mixture of the six amino acids missing from SNM, as well as on TSA for growth control. The screening was performed in three biological replicates, each with at least two technical replicates, resulting in a total of 757 pinned colonies. After two days of incubation at 37°C, plates were imaged using a semi-automatic imager (spImager, S&P Robotics). Colony size was measured using IRIS^47^ in the default mode “Colony growth”, which was also used as a growth metric. Data quality and reproducibility were assessed by visual inspection of the IRIS output and replicate correlation. To account for enhanced growth of colonies in the outer frame of each plate, colony growth was normalized based on the average increase observed in outer frame colonies.

Colony size comparisons were then conducted to assess the effects of amino acid supplementation, as outlined in **Fig. S11C**. Briefly, isolates forming colonies of equal or smaller size than the genetic tyrosine auxotrophs (*S. aureus* LS1^48^, *tyrA* mutants of USA300 JE2 and Newman) on SNM were classified as having a growth defect. Isolates that formed larger colonies on SNM+TYR or SNM+6AA compared to SNM were identified as stimulated by the corresponding amino acid(s). Isolates that exhibited similar growth across all three tested conditions were considered unaffected by amino acid supplementation.

### Nasal colonization of gnotobiotic mice

For *in vivo* experiments of nasal colonization, bacterial inocula containing either *S. aureus* alone or a mixture of *S. aureus* and a SynCom were prepared in PBS-glycerol and stored at -80°C until use. Gnotobiotic C57BL/6N mice were anesthetized and 5 to 10 µL of the bacterial inoculum was distributed on both nostrils, with a target dose of 2×10^4^ CFU of *S. aureus* and 3×10^5^ CFU of the SynCom per nose. Seven days after inoculation, mice were sacrificed to surgically harvest nose, feces, lung and liver. All samples were vortexed for one minute in sterile PBS prior to plating on TSA+Blood or Chromagar *S. aureus* plates to evaluate *S. aureus* colonization (nose, feces) and the absence of systemic dissemination (lung, liver). All animal experiments were conducted in accordance with German laws after approval (IMIT 02/21G) by the local authorities (Regierungspraesidium Tuebingen).

### DNA extraction, amplification and sequencing

For 16S rRNA gene amplicon sequencing, metagenomic DNA was extracted from nasal swabs using the HostZero Microbial DNA kit (Zymo Research). DNA amplification, amplicon library pooling, and V1-V3 16S rRNA gene amplicon sequencing were conducted by the NGS Competence Center Tübingen (NCCT), as described previously^49, 50^. The complete 16S rRNA gene amplicon sequencing results are detailed in **Table S1**, and the raw data will be made publicly available on NCBI upon publication. The DNA extraction and the whole-genome sequencing (WGS) of individual strains were performed by the NCCT or by the Center for Biotechnology Bielefeld (CeBiTec), using Illumina NovaSeq and/or PromethION devices **(Supplementary Methods) (Table S3)**. Circular genomes are available on the BioProjects PRJNA1199961 (newly identified *Corynebacterium* species) and PRJNA1247435 (other species).

### Genomic analyses

16S rRNA gene amplicon sequencing data were analyzed as described previously^50^. For individual strains, sequencing reads were assembled into circular genomes using Flye^51^, Canu^52^ or Unicycler^53^ depending on the sample **(Supplementary Methods)**. These genomes were then classified using pubMLST (https://pubmlst.org/), which compares MLST profiles of the sample to a proprietary database, and using the Type Strain Genome Server TYGS^20^ (https://tygs.dsmz.de), which calculates digital DNA-DNA hybridization (dDDH) values between two genomes and compares them. Genome pairs with a dDDH value over 99.5% were considered of clonal origin. Genome annotation was performed using Bakta^54^ (https://github.com/oschwengers/bakta) or Prokka^55^ (https://github.com/tseemann/prokka).

The frequency of frame-shift mutations in the *tyrA* gene across *S. aureus* isolates was determined as follows. *S. aureus* WGS data was retrieved from Refseq (n=9340). The DNA sequence of *tyrA* from *S. aureu*s Newman (NWMN_1277) was used as a query in a BLAST^56^ search against obtained WGS data featuring the assembly level “Chromosome”, “Complete Genome” or “Contig”. Identified loci were aligned against the query sequence using Clustal Omega provided in the msa R package^57^ and the TyrA ORF was determined. Truncated versions of TyrA featuring less than 363 amino acids were classified as disrupted.

### Statistical analyses

Statistical analyses were performed using GraphPad (v9.3.1 and v10.2.1) or the R stats package. Data normality was assessed using the Shapiro-Wilk test. Differences between group means were assessed using Analysis of Variance (ANOVA) followed by Dunnett’s, Tukey’s or Šídák’s post-test comparisons, as specified in the corresponding figures. For the high-throughput screening of tyrosine auxotrophs, differences between mean colony sizes were evaluated using a t-test followed by Bonferroni correction to control for multiple comparisons. The medians of bacterial concentrations recovered from mice noses and feces were compared using the Kruskal-Wallis test with Dunn’s correction. Adjusted *P* values below 0.05 were considered significant and displayed as \**P*_adj_<0.05, \*\**P*_adj_<0.01, \*\*\**P*_adj_<0.001, \*\*\*\**P*_adj_<0.0001.

## Acknowledgments

We acknowledge the volunteers who participated in this study. We thank A. Peschel and B. Krismer for their support and helpful discussions, as well as L. Lo Presti for reviewing the manuscript and providing critical comments. Special thanks to J. Rudkin (Glasgow), J. Larsen (Copenhagen) and A. Schmidt (Tübingen) for providing a large collection of *S. aureus* strains. We acknowledge A. Weber, L. Maier and the animal keepers for their support in the gnotobiotic facility. We thank A. Bitzer and R. Stemmler for their help in establishing and managing the strain collection; C. Rückert-Reed and T. Busche (Bielefeld) for their support with genome sequencing and analysis; K. Wesner and L. Michaelis for their assistance during animal experiments; the team of H. Brötz-Oesterhelt for providing their camera system; R. van Dalen and A. Kengmo Tchoupa for their valuable advice on metagenomic analysis and cloning. NGS sequencing methods were performed with the support of the DFG-funded NGS Competence Center Tübingen (INST 37/1049-1) and the Institute for Medical Microbiology, University Hospital Tübingen. Finally, we acknowledge D. Kraus, S. Schultz and G. Bauer-Haffter for managing the project funding. This work was funded by the Deutsche Forschungsgemeinschaft (DFG, German Research Foundation) under Germanýs Excellence Strategy – EXC 2124 – 390838134 (L.C., L.T.A., H.L., S.He.) and the German Centre for Infection Research (DZIF) (S.He.). L.C. was also supported by Athene Grant from the Gender Equity Office of the University of Tübingen. The funders had no involvement in the study design, data collection and analysis, decision to publish, or manuscript preparation.

## Author information

### Contributions

L.C. and S.He. conceptualized the study. L.C. designed experiments and performed most of the data visualization. L.C., J.F., D.G., A.L., J.M., M.N.D., J.R., K.B. and S.Ha. conducted and analyzed experiments. D.G., J.J.P., M.O., L.J.K. performed the bioinformatic analysis. V.S., J.K., L.T.A. and H.L. contributed material and analysis tools. L.C. drafted the manuscript. L.C., L.T.A., H.L. and S.He. acquired funding. All authors reviewed and approved the paper.

### Corresponding authors

Correspondence to Laura Camus and Simon Heilbronner. Requests for resources should be directed to and will be fulfilled by the lead contact, Simon Heilbronner.

## Supplementary information

**Supplementary Document A.** PDF file containing Supplementary Methods and Supplementary Figures S1-S12.

**Supplementary Document B.** Excel file containing Supplementary Tables S1-S7.

