## Supplementary Document A for "Tyrosine availability shapes *Staphylococcus aureus* nasal colonization and interactions with commensal communities"

### Supplementary information

**Supplementary methods** – Pages 2 to 4.
**Supplementary figures** – Pages 5 to 17.
**Supplementary tables** – Separated Excel file. Table list and legends on page 18.
**Supplementary references** – Page 19.

### Supplementary methods

#### Isolation and characterization of nasal communities

For each volunteer, a total of seven nasal swabs were collected over three consecutive days to conduct metagenomic sequencing (six swabs) and strain isolation (one swab).

***Metagenomic sequencing.*** Six swabs were individually subjected to metagenomic DNA extraction using the HostZero Microbial DNA kit (Zymo Research) right after sampling. The resulting DNA was pooled prior to V1-V3 16S rRNA gene metagenomic sequencing as described elsewhere[1,2]. For this, the genomic DNA was quantified with a Qubit dsDNA BR/HS Assay Kit (Thermo Fisher) and normalized to a maximum of 5 ng Input for library preparation. The first step PCR was performed in 25 µl reactions including KAPA HiFi HotStart ReadyMix (Roche), custom primers and template DNA (PCR program: 95°C for 3 min, 30x (98°C for 20 sec, 69°C for 15 sec, 72° for 15 sec), 72°C for 5 min). First PCR products were purified using 28µl AMPure XP beads and eluted in 50µL 10mM Tris-HCl. Indexing was performed in the second step PCR including KAPA HiFi HotStart ReadyMix (Roche), index primer mix (Illumina Nextera XT Index Kit v2), purified first PCR product as template (PCR program: 95°C for 3 min, 8x (95°C for 30 sec, 55°C for 30 sec, 72°C for 30 sec), 72°C for 5 min). After another bead purification (20µl AMPure XP beads, eluted in 30µL 10mM Tris-HCl) the libraries were checked for correct fragment length on an agarose gel, quantified with a Qubit dsDNA BR Assay Kit (Thermo Fisher) and pooled equimolarly. The pool was sequenced on an Illumina MiSeq device with a v2 sequencing kit with 101,8,8,401 read length.

***Strain isolation and first identification.*** The seventh nasal swab was subjected to strain isolation procedure as described previously[3]. A first fraction of the nasal swab was plated on 4 different rich agar media: Columbia agar with 5% sheep blood (Fisher Scientific), Schaedler agar with 5% sheep blood, Chocolate agar with 10% sheep blood, Schaedler agar supplied with 5% horse blood, kanamycin, and vancomycin (Universitätklinikum Tübingen). Plates were incubated for 3 to 10 days under aerobic, anaerobic or 5% $CO_2$ conditions. In parallel, the second fraction of the swab was used to inoculate two liquid broths: BHI (Roth) supplemented with 0.2% Tween 80 (Sigma) for aerobic growth during 12 h, and Thioglycollate medium (Sigma) for anaerobic growth during 48 h. The resulting cultures were then plated on Columbia agar with 5% sheep blood or on Schaedler agar with 5% sheep blood and incubated for 2 to 7 days in aerobiosis or anaerobiosis. For each agar plate, the colony morphotypes were differentiated by visual inspection at the end of the incubation period, and the bacterial concentration of each morphotype was determined by colony enumeration. Two colonies per morphotype were picked and reisolated on the appropriate agar medium prior to species identification using matrix-assisted laser desorption/ionization-time-of-flight mass spectrometry (MALDI-TOF MS) as described elsewhere[4]. To promote cell lysis and accurate species identification, 1µL of 70% formic acid was added to the smeared colonies on the MS target plate prior addition of the MS matrix (2). The analysis was then performed by the Institute of Medical Microbiology and Hygiene (University Hospital Tübingen, Germany) using the Bruker Daltonik MALDI Biotyper and a composite database of 9128 Main Spectrum Profiles (Bruker Daltonik Database, Filamentous Fungi, Clinical SR database, Database Tübingen). Species identification was considered explicit when the MS score exceeded or was equal to two. Two to four isolates per species and per volunteer were stored to constitute the LaCa collection.

#### Whole genome sequencing of individual isolates

***Nanopore sequencing.*** Nanopore sequencing was performed by the Center for Biotechnology (CeBiTec Bielefeld) on a subset of strains **(Table S3)**. Genomic DNA was extracted from cell pellets using the Macherey-Nagel NucleoSpin Microbial DNA Kit. Sequencing libraries were prepared with the Native Barcoding Kit (SQK-NBD114.96, Oxford Nanopore). The DNA repair and end-prep steps were performed with the recommended reagent volumes for samples with DNA concentrations between 8ng/µL and 18 ng/µL. For samples with DNA concentrations below 8 ng/µL, the reagent volumes were doubled, and for samples with concentrations above 18 ng/µL, the volumes were halved. Barcoding was performed using equal amounts of end-prepped DNA, blunt/TA ligase mix, and EDTA. Sequencing was carried out using FLO-PRO114M flow cells on a PromethION P2 Solo device. Base calling was performed with Guppy base caller v6.3.9.

***Hybrid sequencing.*** Hybrid sequencing was conducted by the NGS Competence Center Tübingen (NCCT) on a subset of strains **(Table S3)**. For bacterial lysis, a cell pellet was resuspended in 600 µL ATL buffer (Qiagen #939011) and transferred to a ZR BashingBead Lysis Tube (Zymo Research #S6012-50). The tube was vortexed horizontally for two minutes on a vortex shaker. To optimize the DNA extraction, the supernatant was taken off and digested with RNAse A (Qiagen). The DNA was then automatically purified with the QIAamp 96 QIAcube HT kit (Qiagen #51331) with additional proteinase K on a QIAcube HT following the manufacturers' instructions. The genomic DNA was quantified with a Qubit dsDNA BR Assay Kit (Thermo Fisher) and DNA integrity was checked by agarose gel electrophoresis. Libraries for short-read sequencing were prepared using the Illumina DNA Prep (M) Tagmentation kit according to the manufacturer's protocol with 500 ng DNA input and 5 cycles indexing PCR. Libraries were checked for correct fragment length on an Agilent 2100 BioAnalyzer and pooled equimolarly. The pool was sequenced on an Illumina NovaSeq device with a NovaSeq 6000 SP v1 sequencing kit with 2 x 150 bp read length. The sequencing was demultiplexed with bcl2fastq (v2.19.0.316) and quality checked with fastq (v0.20.1) and visualized with MultiQC (v1.7). For long-read sequencing the Ligation Sequencing Kit (LSK) 109 (Oxford Nanopore) was used with native barcoding, following the manufacturer's protocol, with 500 ng DNA input per sample. Library size was assessed on a FEMTO Pulse (Agilent) and libraries were pooled equimolarly before sequencing on a FLO-PRO002 flow cell on a Nanopore PromethION device. Base calling was performed in high-accuracy mode with Guppy base caller v4.0.11 and with MinKNOW operating software v20.06.18.

***Genome assembly and annotation.*** Long read sequencing reads were assembled into circular genomes using Flye[5] and annotated with Bakta[6]. Hybrid sequencing were assembled using the nf-core pipeline bacass (https://nf-co.re/bacass/2.0.0/), which includes Unicycler[7] and Canu[8] for assembly into circular genomes and Prokka[9] for annotation.

### Supplementary figures

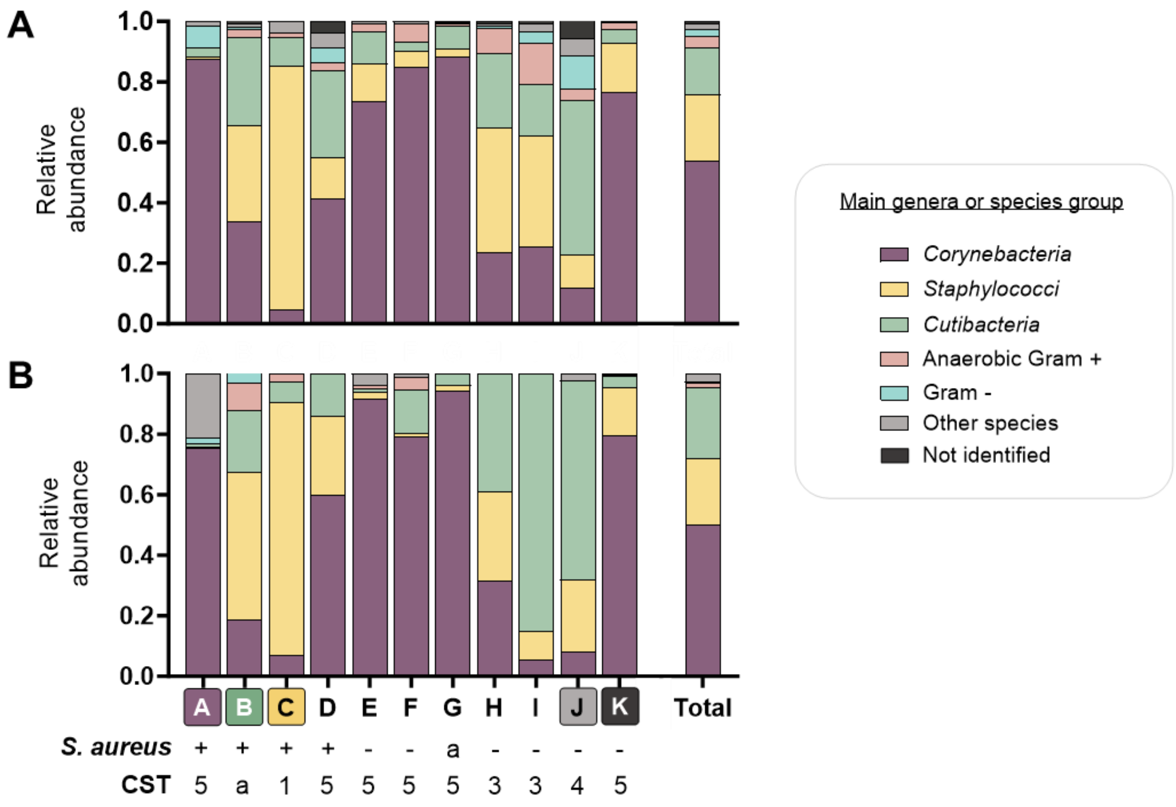


**Fig. S1: Composition and structure of 11 nasal communities, at the group level.** Relative species abundance determined by 16S rRNA gene sequencing of nasal metagenomic DNA (**A**) or by isolation and enumeration of nasal commensals on plates (**B**), from the nasal microbiomes of eleven healthy volunteers (A-K). Each color designates the genus or the group of the 20 most abundant species across all volunteers. Less abundant species were assigned to "Other species". Species with a missing or unclear identification were assigned to "Not identified". Below the bars, "+" indicates the volunteers in which *S. aureus* was detected by both techniques, and numbers correspond to the community state type (CST) assigned to the corresponding community. "a" refers to ambiguous *S. aureus* carriage (detected by only one method) or CST classification (balanced proportion of several species). Each bar displays the results of one replicate.

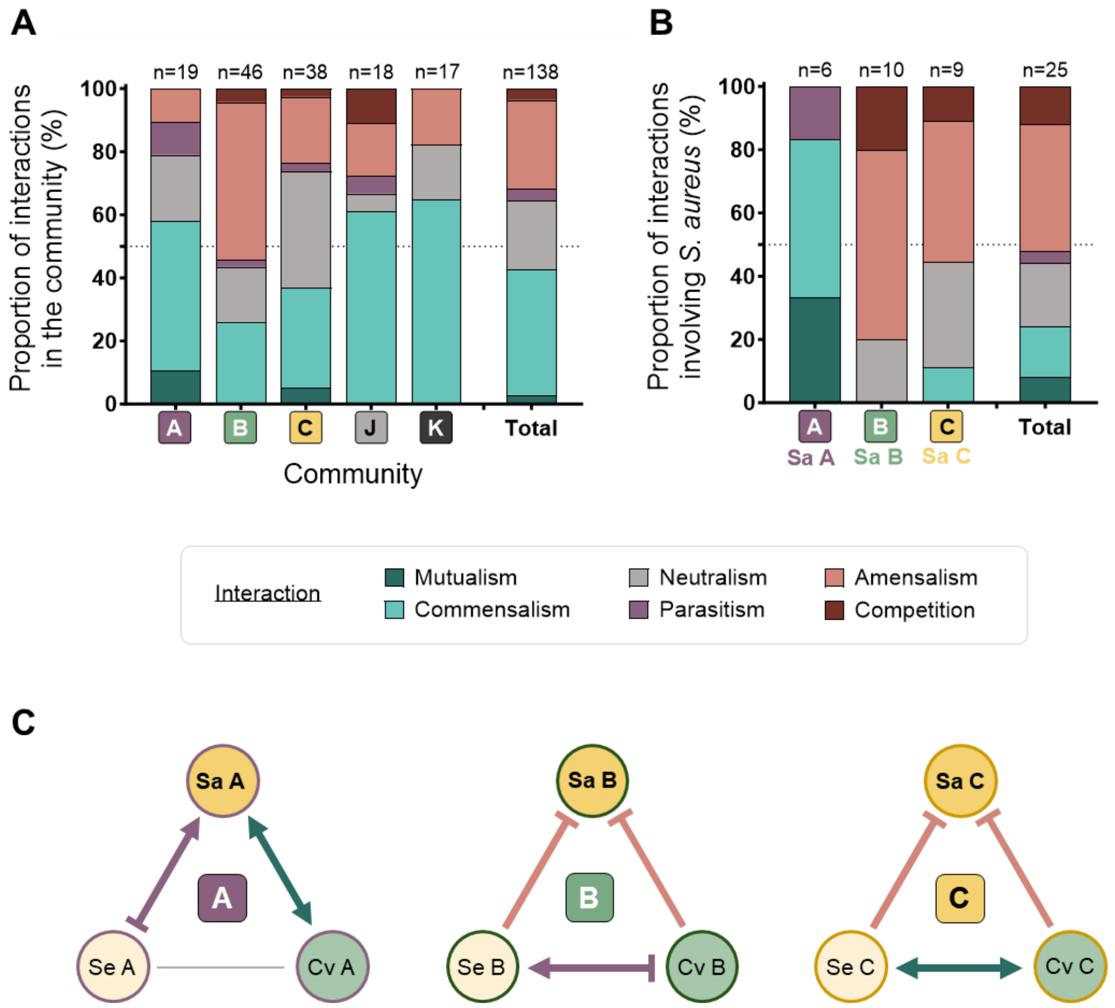


**Fig. S2: Binary interactions in the nasal communities A, B, C, J and K.**
For each community, interactions were determined for every pair of nasal commensals on SNM agar and using colony area as a growth metric. Changes in the colony area were used to define the type and the strength of the interaction.
**A, B.** The interactions between all commensals (**A**) or involving *S. aureus* (**B**) were counted to plot their proportion within each community. The total number of interactions determined within each community is marked on the top of the bar.
**C.** Binary interactions between the strains of *S. aureus* (Sa), *S. epidermidis* (Se) and *C. avidum* (Cv) isolated from the communities A, B and C. Interactions were categorized as in the panels A and B.

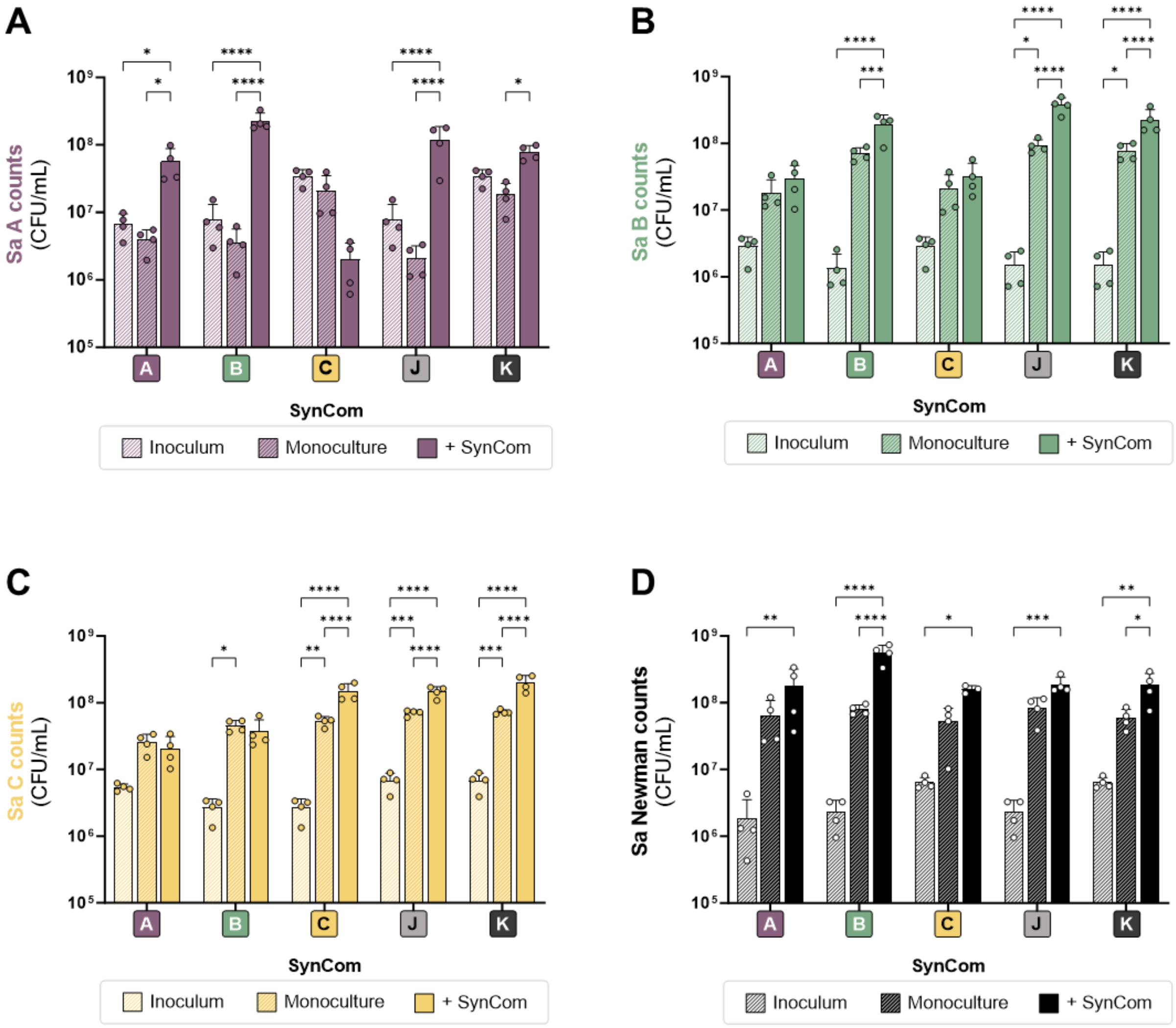


**Fig. S3: Impact of SynComs on *S. aureus* proliferation in SNM.**
Barplots depicting the bacterial counts of *S. aureus* in the inoculum, and after incubation for 3 days in monoculture and in synthetic culture with five different SynComs (A, B, C, J, K) on SNM agar. Each bar displays the mean + SD of four biological replicates. Statistical significance is indicated by $*P_{adj}<0.05$, $**P_{adj}<0.01$, $***P_{adj}<0.001$, $****P_{adj}<0.0001$ (Two-way ANOVA with Tukey's correction). Panels display the results for Sa A (**A**), Sa B (**B**), Sa C (**C**) and Sa Newman (**D**).

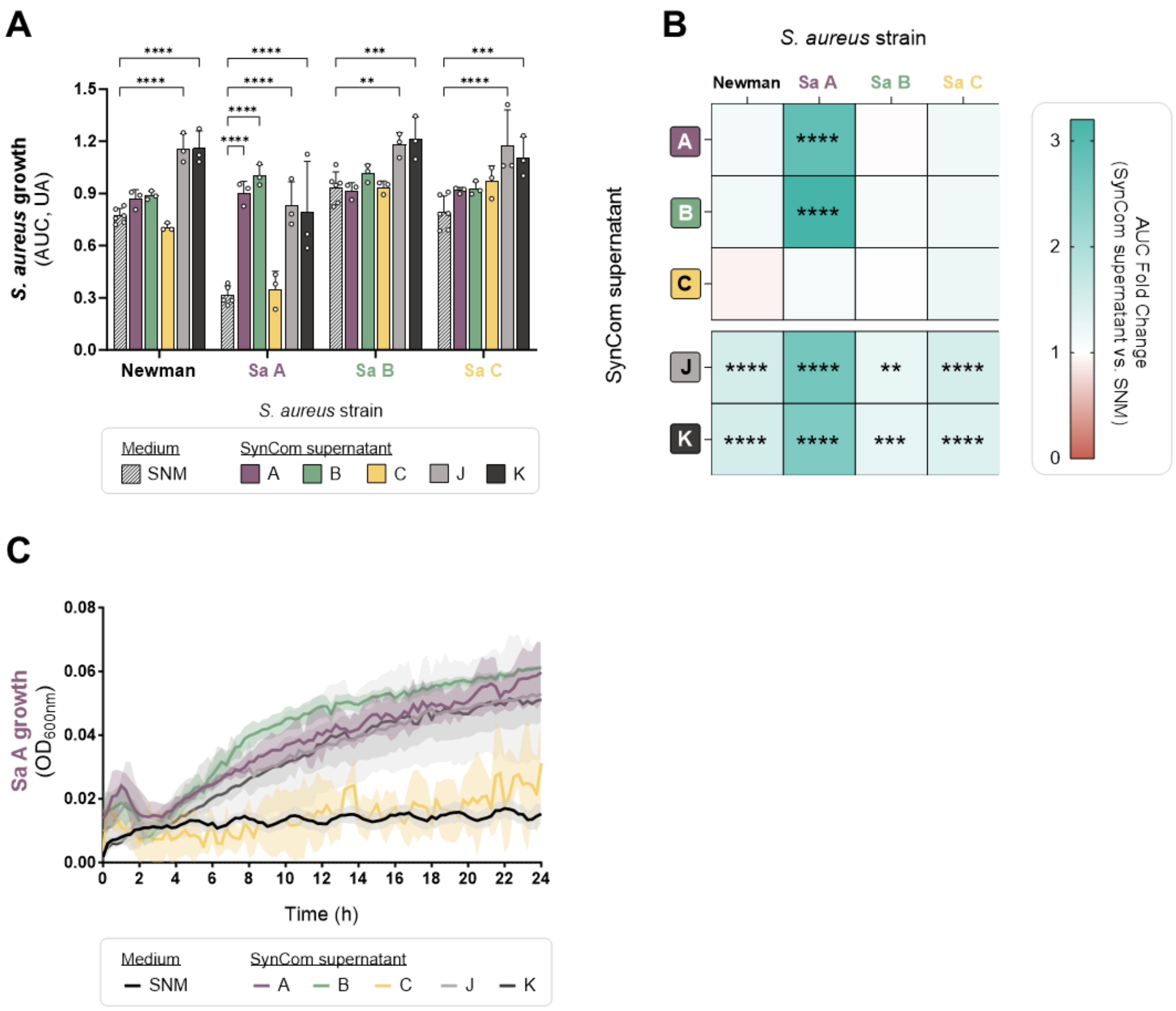


**Fig. S4: Impact of SynCom supernatant on *S. aureus* proliferation in SNM.**
**A.** Barplots showing the bacterial growth of four *S. aureus* strains cultivated in SNM or in the presence of 50% concentrated spent culture supernatant of the SynComs A, B, C, J and K. Each bar displays the mean AUC + SD from 24h growth kinetics.
**B.** Heatmap showing the AUC fold-change induced by the supernatant of the five SynComs in SNM, for four *S. aureus* strains. Positive fold-changes (blue) and negative fold-changes (red) indicate increases or decreases in *S. aureus* growth with the supernatant relative to the SNM control, respectively.
**C.** Complete growth kinetic of Sa A in SNM or in the presence of the supernatant of the five SynComs. Lines show the mean ± SD of $OD_{600nm}$ measurements, taken every 15 minutes over 24 hours.
**For all panels:** The data is based on at least three biological replicates of $OD_{600nm}$ growth kinetics. Statistical significance is indicated by *$P_{adj}<0.05$, **$P_{adj}<0.01$, ***$P_{adj}<0.001$, ****$P_{adj}<0.0001$ (Two-way ANOVA with Dunnett's correction).

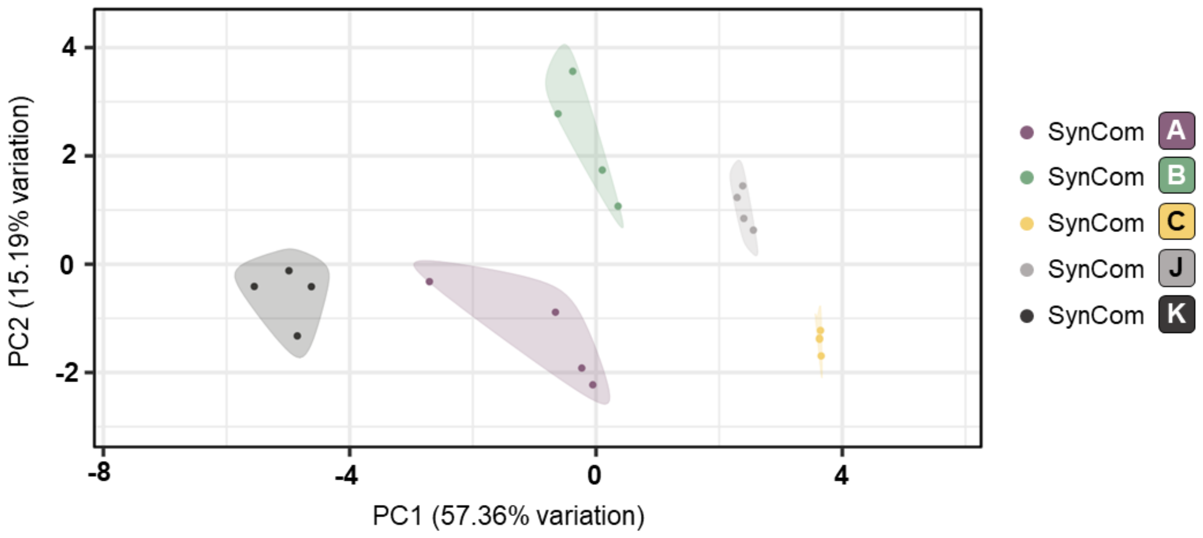


**Fig. S5: Amino acid profile of the 5 SynComs cultivated in SNM.**
Principal component analysis based on 17 amino acids detected in the extracts of the five SynComs cultivated for 3 days on SNM. Each dot is based on the LC-MS data from one biological replicates (four per SynCom). Ellipses represent the 95% confidence interval of each group. Group differences and dispersion were assessed using PERMANOVA (*P* = 0.001, $R^2$ = 0.8974), and PERMDISP (*P* = 0.005).

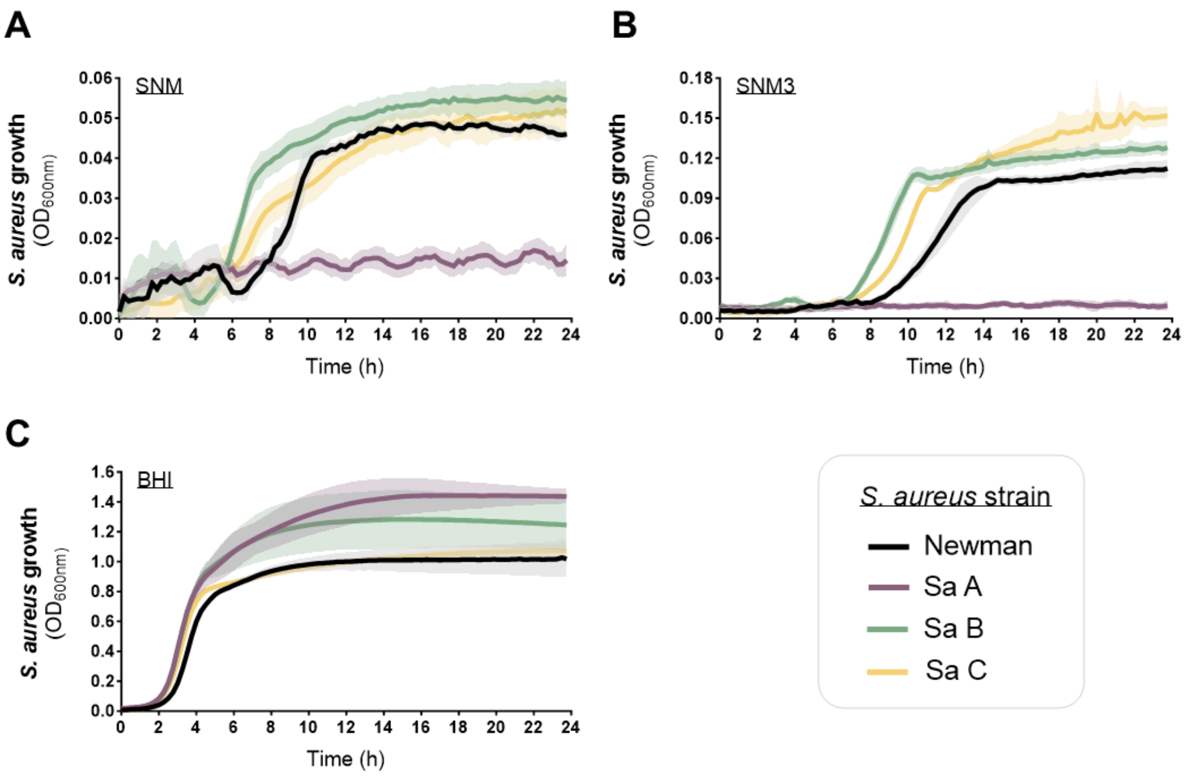


**Fig. S6: Proliferation of four *S. aureus* strains in various media.**
Growth kinetics in SNM (**A**), SNM3 (**B**) or BHI (**C**) for the *S. aureus* strains Newman, Sa A, Sa B or Sa C. Lines represent the mean ± SD of at least three biological replicates of $OD_{600nm}$ measurements, taken every 15 minutes over 24 hours.

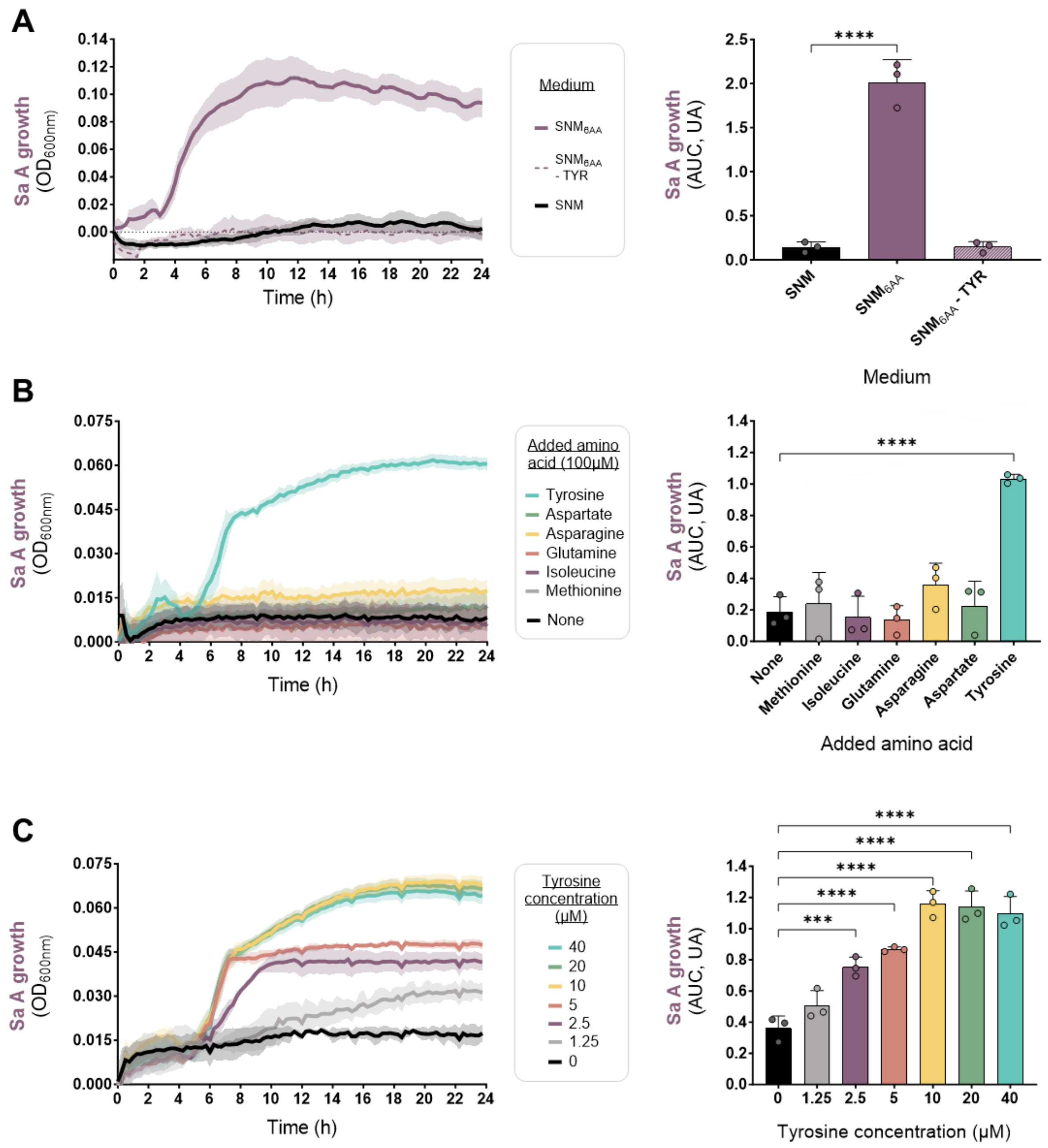


**Fig. S7: Impact of amino acid supply on the proliferation of *S. aureus* Sa A in SNM.**
**For all panels:** growth kinetics are displayed on the left and corresponding AUC bar plots are shown on the right. Lines or bars represent the mean ± SD of three biological replicates of $OD_{600nm}$, taken every 15 min over 24 hours. Statistical significance is indicated by $***P_{adj}<0.001$, $****P_{adj}<0.0001$ (ANOVA with Dunnett's correction).
**A.** Proliferation of Sa A in SNM, in SNM supplemented with a mixture of the six amino acids missing from SNM (6AA), or with the same mixture from which tyrosine was removed (6AA-TYR). 100µM of each amino acid were used.
**B.** Proliferation of Sa A in SNM, or in SNM supplemented with 100µM of each individual amino acid missing from SNM.
**C.** Proliferation of Sa A in SNM, or in SNM supplemented with increasing concentrations of tyrosine.

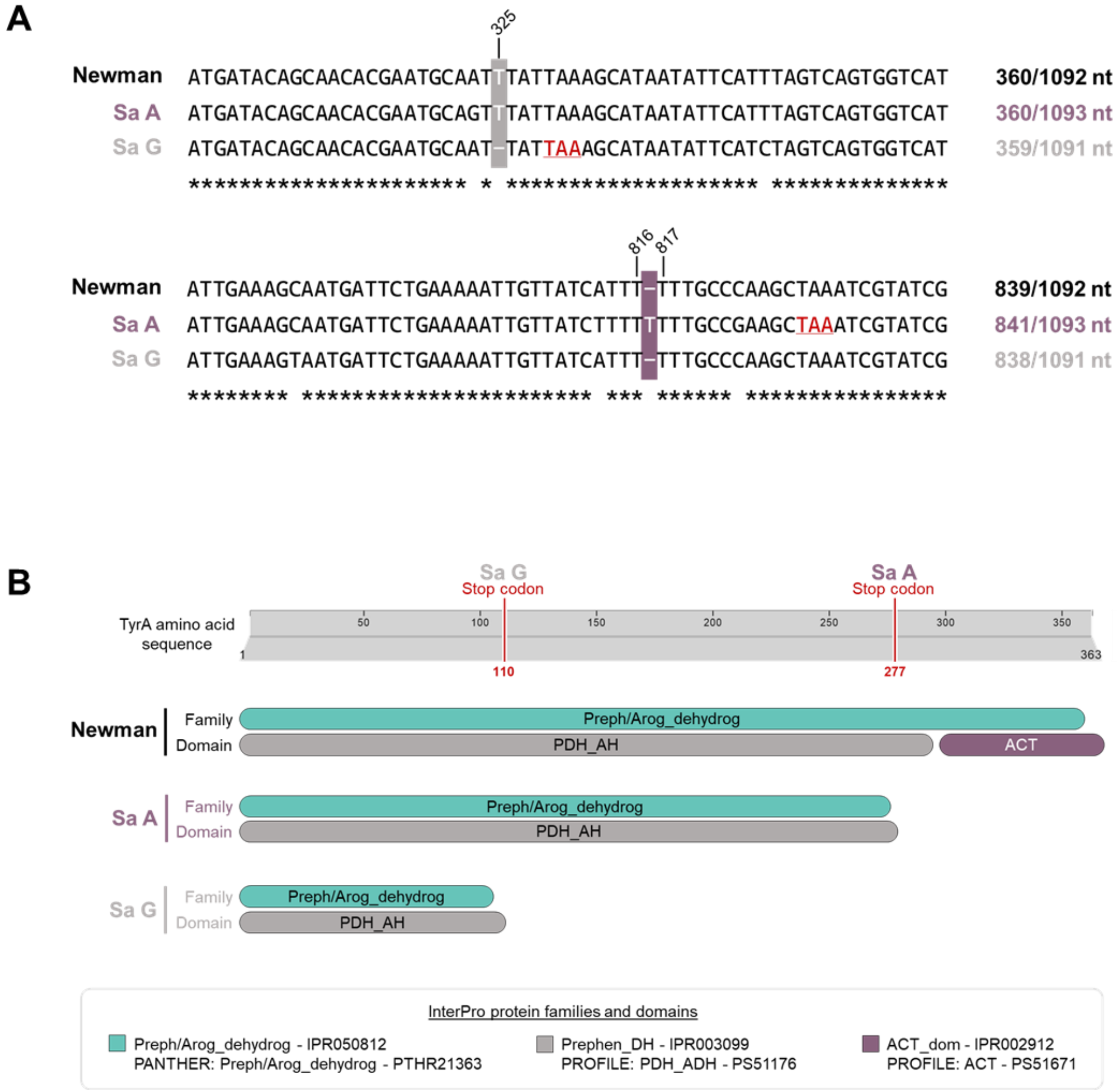


**Fig. S8: Identification of deleterious mutations in the *tyrA* gene of Sa A and Sa G.**
**A.** Sequence alignment of the *tyrA* gene *S. aureus* Newman, Sa A and Sa G. The T insertion and deletion events are highlighted in purple and gray, respectively, with the resulting stop-codon marked in red. Nucleotide positions of these mutations relative to Newman sequence are shown at the top. Adapted from Clustal Omega multiple sequence alignments.
**B.** Protein family and domain prediction of TyrA from *S. aureus* Newman, Sa A and Sa G. The top scale shows the full amino acid sequence of TyrA and highlights the positions of the stop codons identified in Sa A and Sa G. Adapted from InterPro. PDH_AH: prephenate/arogenate dehydrogenase. ACT: domain found in aspartate kinase, chorismate mutase and TyrA.

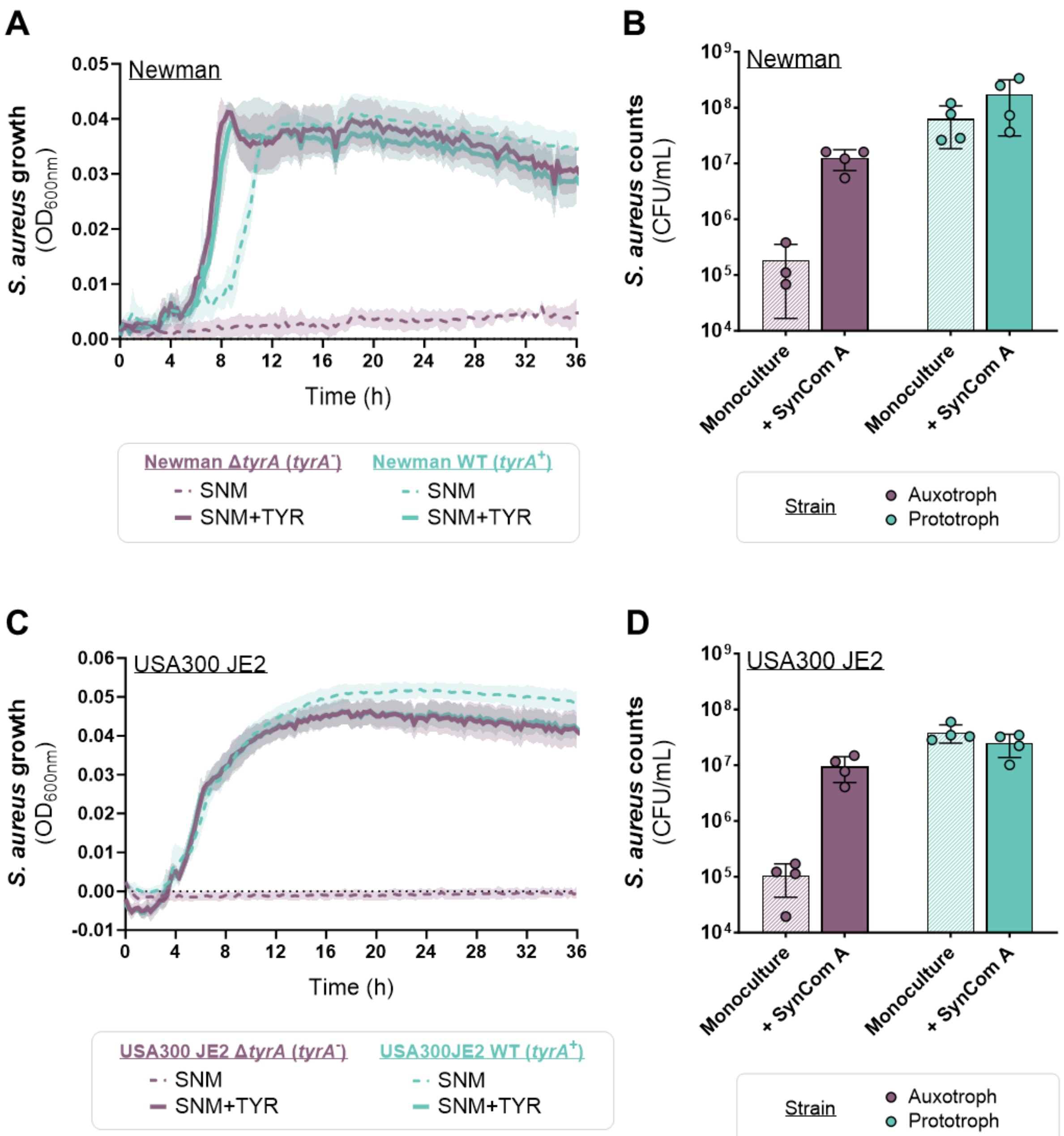


**Fig. S9: Effect of *tyrA* deletion on the proliferation of *S. aureus* Newman and USA300 JE2 in SNM.**
For all panels, the *S. aureus* strains Newman (**A, B**) and USA300 JE2 (**C, D**) with an impaired *tyrA* gene (Δ*tyrA* strains) are referred as *tyrA*$^-$ and shown in purple, while the strain with native *tyrA* gene (WT strains) are named *tyrA*$^+$ and shown in blue.
**A, C.** Growth kinetics in SNM and SNM + tyrosine (TYR). Lines represent the mean ± SD of four biological replicates of $OD_{600nm}$ measurements, taken every 15 minutes over 36 hours.
**B, D.** Barplots showing the bacterial counts of *S. aureus* cultivated on SNM agar in isolation or in the presence of the SynCom A over three days. Bars represent the mean ± SD of four biological replicates. Statistical significance was assessed using two-way ANOVA with Šídák's correction.

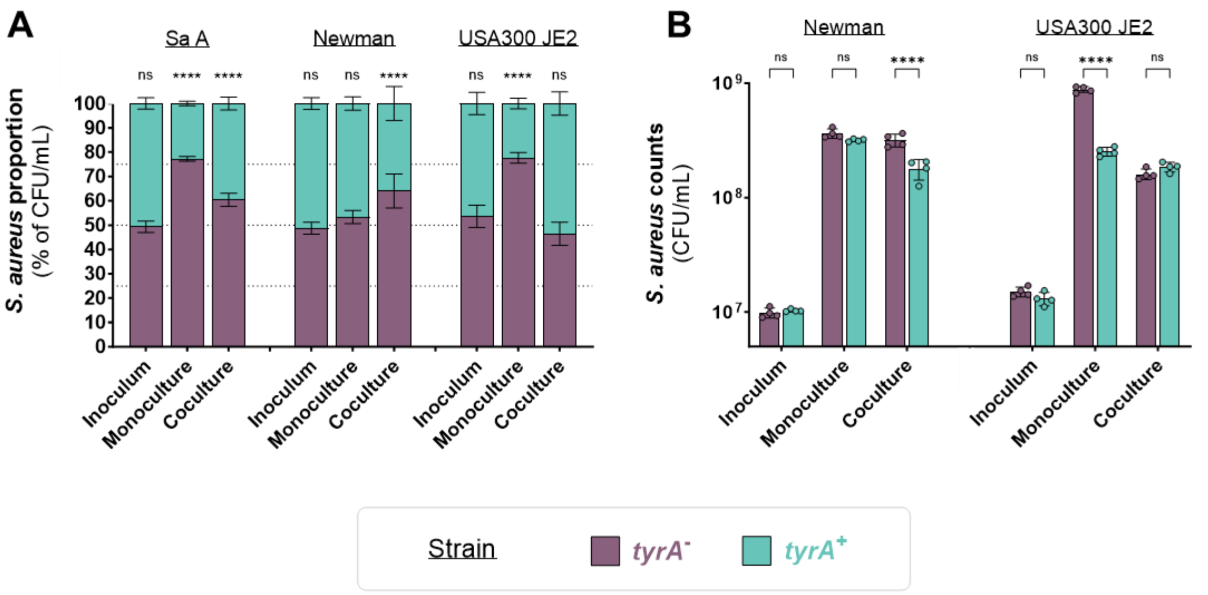


**Fig. S10:** **Effect of *tyrA* deficiency on *S. aureus* fitness in SNM.**
Barplots showing the bacterial proportion (**A**) or counts (**B**) of *tyrA*$^+$ and *tyrA*$^-$ strains of *S. aureus* in the inoculum, cultivated in SNM + tyrosine either in isolation or cocultured with each other over 24 hours. For all panels, the *S. aureus* strains with an impaired *tyrA* gene (Sa A WT, Newman and USA300 JE2 Δ*tyrA*) are referred as *tyrA*$^-$ and shown in purple, while the strains with a restored or native *tyrA* gene (Sa A-*tyrA*, Newman and USA300 JE2 WT) are named *tyrA*$^+$ and shown in blue. Dashed lines indicate the proportions 25%, 50% and 75% on the panel A. Bars represent the mean ± SD of four biological replicates. Statistical significance is indicated by ****$P_{adj}$<0.0001 (Two-way ANOVA with Šídák's correction).

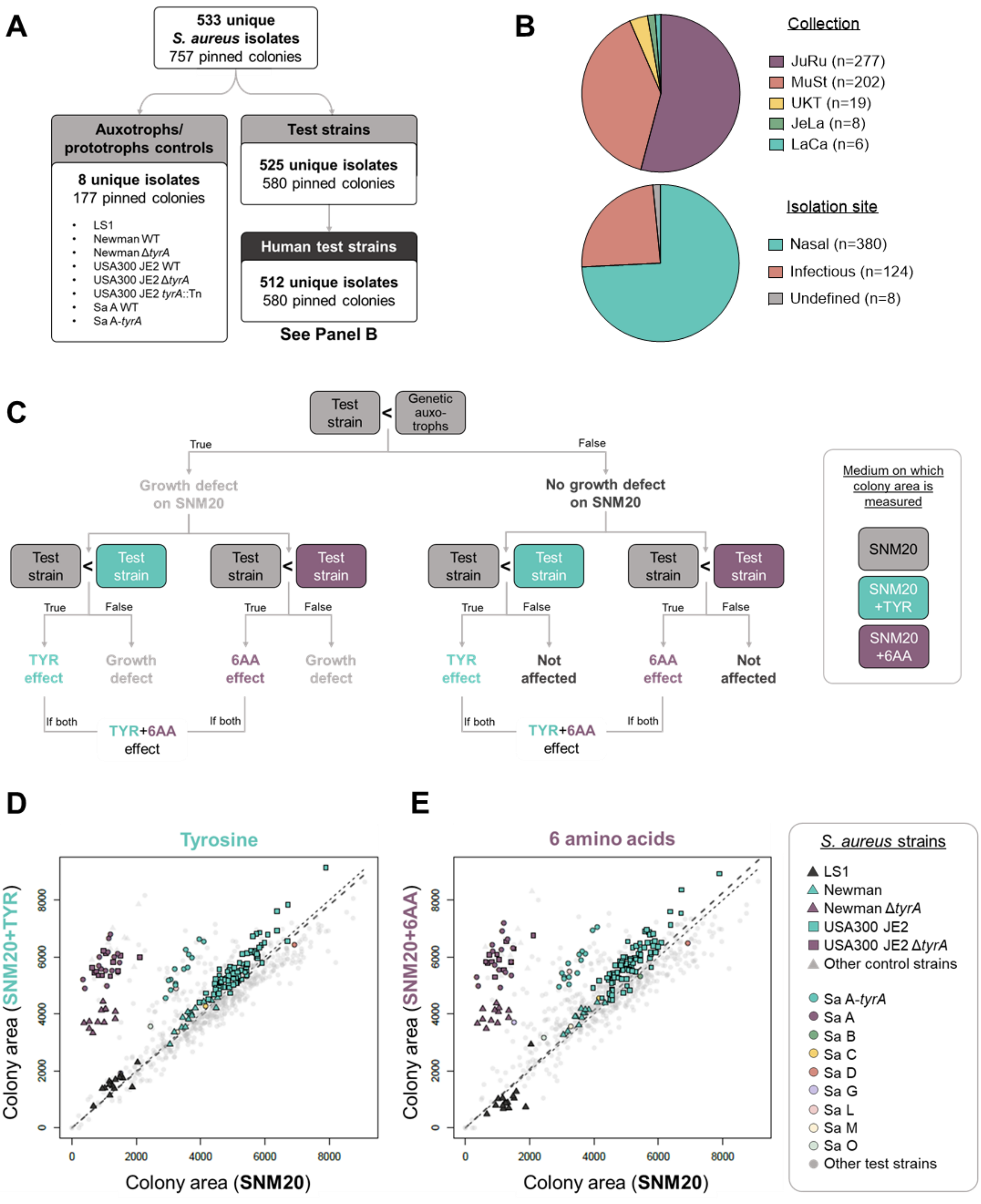


**Fig. S11: Screening for phenotypic tyrosine auxotrophies in *S. aureus*.**

**A.** Numbers of unique isolates and corresponding pinned colonies screened for phenotypic auxotrophy.

**B.** Pie charts illustrating the collections from which the 512 human isolates were obtained (top) and their isolation sites (bottom).

**C.** Decision tree used to analyze the growth phenotypes of *S. aureus* isolates pinned on SNM20 and SNM20 supplemented with tyrosine (TYR), or the six amino acid mixture (6AA). Genetic auxotrophs used as a negative controls to identify isolates with a growth defect on SNM20 include *S. aureus* strains LS1 and *tyrA* mutants of Newman and USA300 JE2.

**D, E.** Dotplots showing the relationship between growth on SNM20 and SNM20 supplemented with tyrosine (**D**), or the six amino acid mixture (**E**), for the complete collection of *S. aureus* isolates (n= 757 depicted colonies; n=533 unique isolates). Dots represent the colony area from one biological replicate. Controls and strains of interest are shown in different shapes and colors. The diagonal dashed lines represent the linear regression corresponding to a ratio of one (thin line) or the actual linear regression (thick line).

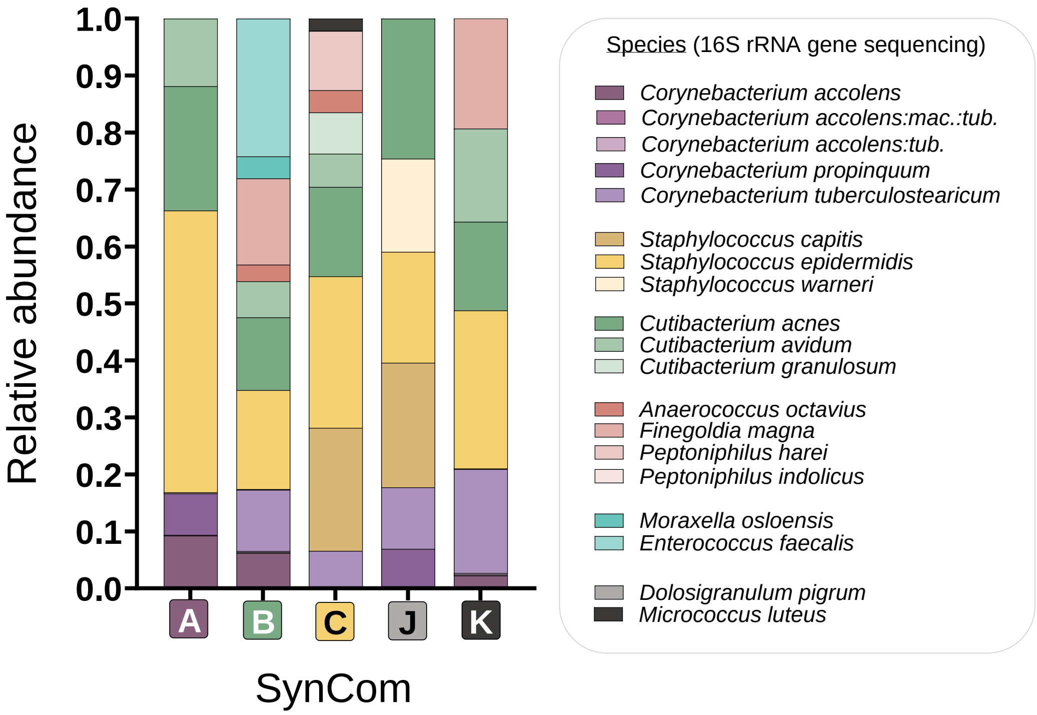


**Fig. S12: Species composition and structure of the SynComs A, B, C, J and K.** Relative species abundance in frozen SynComs stocks determined by 16S rRNA gene sequencing. Each color designates a species. Each bar displays the results of one replicate.

### Supplementary tables (separated Excel file)

**Table S1: Complete 16S rRNA gene sequencing results.**
Absolute and relative abundance of all Amplicon Sequence Variants (ASV) determined by 16S rRNA gene sequencing of nasal metagenomic DNA from eleven healthy volunteers (A-K). Each value was obtained from one replicate.

**Table S2: Summary of 16S rRNA gene sequencing and strain isolation results.**
Number of ASV, isolates or species identified by the two approaches for the eleven volunteers (A-K). Community State Types[10] (CST) were assigned according to the most abundant species identified by 16S rRNA gene sequencing.

**Table S3: LaCa nasal strain collection.**
Strains isolated from the nasal swabs of eleven healthy volunteers (A-K). For each strain, species was first identified via MALDI-MS and the abundance within the swab was determined by enumeration on agar plates. The genome of each strain was then sequenced to confirm the species identification and determine the intraspecies clonality. “LaCa identifier” designates the strain number in the collection LaCa, whereas “Special identifier” indicates the strain name used in this manuscript.

**Table S4: Amino acid relative concentrations detected by LC-MS in five SynComs.**
Ratio of $^{12}C$ signal of the sample and $^{13}C$ signal of the internal standard, for 17 amino acids detected by LC-MS analysis on SynCom cultures (A, B, C, J, K). Each value was obtained from one biological replicate (four per SynCom). Optical density measurements used to normalize the data to SynCom growth are also presented.

**Table S5: Non-LaCa bacterial strains used in this study.**
*E. coli* and *S. aureus* wild-type and mutant strains, as well as *S. aureus* isolates from the collections of Munich students (MuSt), Jesper Larsen (JeLa), Justine Rudkin (JuRu) or Universitätklinikum Tübingen (UKT). Depending on the collection, strains were identified either by MALDI-MS, isolation on *S. aureus* selective media, or whole-genome sequencing. Associated references: [11–20]

**Table S6: Primers and plasmids used in this study.** Associated references: [12]

**Table S7: Culture conditions of nasal species from the LaCa collection.**
Medium, oxygenation conditions and duration used for cultivating nasal species.
